# Rspo1 dependent WNT signaling safeguards ovarian development by suppressing androgen signaling

**DOI:** 10.64898/2026.09.22.753168

**Authors:** Furong Tang, Natividad Bellido-Carreras, Kheira Bouzid, Aurélie Lardenois, Thomas A. Darde, Magali Dhellemmes, Aurélie Despoux, Mirko Peitzsch, Marie-Christine Birling, Frédéric Chalmel, Marie-Christine Chaboissier, Aitana Perea-Gomez

## Abstract

Embryonic ovarian development requires high levels of WNT/ß-catenin signaling yet the molecular basis of this requirement remains unclear. Mutations in the WNT potentiator RSPO1 impair ovarian development in both mouse and human, leading to gonadal masculinization in XX individuals. Here, we show that RSPO1 acts within a restricted developmental window at the onset of ovarian differentiation and that its absence triggers precocious androgen signaling in the developing ovaries. Moreover, androgen receptor inhibition rescues ovarian development in *Rspo1* mutant mice. Our findings identify RSPO1/WNT/ß-catenin signaling as an essential pathway that suppresses androgen signaling to ensure proper ovarian development in mouse.

## Introduction

In mammals, sexual development is viewed as a three-step process. First, sex chromosomes are inherited at the time of fertilization. Next, gonadal sex determination occurs, whereby an initial undifferentiated gonad adopts a testicular or ovarian fate in XY and XX embryos, respectively. Finally, during sexual differentiation internal and external genitalia adopt a male or female fate depending on the sexual hormones present in the fetus. In XY fetuses, high levels of circulating androgens produced mainly by the testes are required during a restricted time window for the differentiation of male genitalia (including epididymis, vas deferens, seminal vesicles, penis and scrotum) (*1–3*). In addition, testis-derived Anti Mullerian Hormone (AMH) prevents the development of female structures in XY individuals (*4*). In contrast, XX fetuses with low levels of circulating AMH and androgens differentiate female genitalia (including oviducts, uterus, vagina, clitoris and vulva). Tight regulation of androgen signaling is essential for mammalian female sexual development as experimental or pathological increase of androgens results in masculinization of internal and external genitalia of XX fetuses (*2, 3, 5*). However, in contrast to the situation in other vertebrates (*6*), increased androgens levels do not trigger masculinization of the ovary (*7–9*), highlighting that mammalian fetal ovarian development is largely unsensitive to androgen excess and gonadal sex determination occurs independently of sexual hormones (*6, 10*).

Gonadal sex determination occurs at embryonic day 11 (E11) in mice and fifth post-conceptional week (PCW) in humans. In mouse, testis development is initiated by the transient expression of the Y-linked gene *Sry* in XY gonads (*11–13*). SRY dependent up-regulation of SOX9 drives the differentiation of Sertoli cells, the male supporting cells responsible for producing AMH and for orchestrating subsequent steps of testis differentiation (*11, 14–17*). Sertoli cells produce signals that promote the differentiation of interstitial progenitors into steroidogenic fetal Leydig cells (FLC) from E12.5 (*17, 18*). FLCs metabolize cholesterol into androstenedione, which is ultimately converted into testosterone by the HSD17B3 enzyme expressed in Sertoli cells (*19*). Together, fetal Sertoli and Leydig cells coordinate testis differentiation and male sexual development. In XX embryos, the – KTS isoform of WT1 initiates a genetic cascade involving WNT/ß-catenin signaling that drives ovarian development by promoting the differentiation of pre-granulosa cells, the female supporting cells (*20–25*). Pre-granulosa cells formed in successive waves will mature into granulosa cells that surround the oocyte in follicles (*26, 27*). Mature granulosa cells promote the differentiation of stromal progenitors into steroidogenic theca cells (*28*). Theca cells metabolize cholesterol into androgens which are then converted into estrogens by the aromatase enzyme CYP19A1 expressed in granulosa cells (*29*). Therefore, sex steroids production relies on the joined action of supporting and steroidogenic cell types in both testes and ovaries. However, unlike the situation in the testis, steroidogenesis in the ovary does not immediately follow gonadal sex determination. In mice, androgen producing theca cells differentiate after birth, when folliculogenesis is initiated (*30*). In human, folliculogenesis occurs during fetal life and sex steroids are detected from 13-14 PCW in the fetal ovary (*31, 32*).

The secreted protein R-spondin1 (RSPO1) is a potentiator of WNT/ß-catenin signaling essential for ovarian development in human and mouse (*21, 25, 33*). Patients with mutations in RSPO1 exhibit 46,XX testicular or ovotesticular differences in sex development (DSD), with gonads that develop as testes or ovotestes, and varying degrees of genital virilization (*33–36*). In mice, loss of *Rspo1* in XX embryos leads to progressive masculinization of the gonad characterized by precocious maturation of pre-granulosa cells in the fetal ovary, followed by their trans-differentiation into Sertoli-like cells around birth, ultimately resulting in the formation of an ovotestis (*21, 25, 37*). XX *Rspo1* mutants also exhibit abnormal steroidogenic cell differentiation at fetal stages and androgen-dependent masculinization of the internal genitalia resulting in pseudo-hermaphroditism (*21, 25*). Here we investigate the primary causes of sex reversal in XX *Rspo1* mutant gonads in the absence of *Sry*. We first identify the cell populations responsible for abnormal androgen production in the fetal ovaries of these mutants. We then demonstrate that androgens not only masculinize the internal genitalia but also drive ovarian sex reversal in XX *Rspo1* mutants revealing an essential role of RSPO1/WNT/ß-catenin signaling in suppressing both production and signaling of intra-gonadal androgens during mouse ovarian development.

## Results

### Rspo1 function is required at the onset of ovarian differentiation

*Rspo1* is expressed during the entire ovarian development from the undifferentiated gonad stage (E10.5-E11.5) (*33, 38–41*) (Fig. 1, A and B). To determine the critical time window when *Rspo1* is required for ovarian development, we generated a new conditional *Rspo1* floxed allele (*Rspo1^flox^*) allowing CRE mediated deletion of *Rspo1* exon 2 that contains the start codon and signal peptide (fig. S1A). First, we established that deletion of *Rspo1* exon 2 leads to *Rspo1* loss of function. We found that XX *Rspo1^-/-^*gonads where exon 2 of *Rspo1* is constitutively deleted show an ovotestis phenotype, similar to what has been observed with other *Rspo1* loss of function alleles (*21, 25*) (fig. S1, B to F). To address the temporal requirement of *Rspo1* function we used the *Wt1^CreERT2^* mouse line to conditionally delete *Rspo1* in somatic gonadal cells at specific time points (*42*). We found that *Rspo1* is efficiently deleted in the developing gonads of XX *Wt1^CreERT2/+^; Rspo1^flox/flox^* embryos 24 hours after tamoxifen administration to pregnant mothers (Fig. 1C and fig. S2, A to C). When *Rspo1* was deleted at E11.5 (Tamoxifen at E10.5) XX gonads developed as ovotestes and exhibited defects that phenocopy a constitutive *Rspo1* null mutation (*21, 37*) including abnormal down-regulation of the pre-granulosa cell marker FOXL2 (Fig. 1, D and E), precocious maturation of granulosa cells expressing AMH (Fig. 1, H and I), up-regulation of SOX9 in trans-differentiated Sertoli-like cells (Fig. 1, D and E), germ cell loss and abnormal steroidogenic cell differentiation (fig. S2, D, E, H and I) at E17.5. In these mutants, testis-cord like structures were observed at post-natal day 10 (P10) (Fig. 1, L and M). In contrast, when *Rspo1* was deleted from E12.5 (Tamoxifen at E11.5), XX gonads developed as ovaries (Fig. 1, F, J and N, and fig. S2, F and J) with supporting cells differentiating as granulosa cells and no sign of trans-differentiation into Sertoli-like cells. Moreover, deletion of *Rspo1* with *Nr5a1-Cre*, a transgenic line known for driving efficient recombination in gonadal somatic cells from E12.5 (*43, 44*), resulted in ovarian development (fig. S3). These results demonstrate that *Rspo1* function is required in XX somatic gonadal progenitors between E11.5 and E12.5, and that the trans-differentiation of granulosa cells into Sertoli-like cells observed at the end of gestation in XX *Rspo1* mutant ovotestes stems from early defects in critical processes regulated by *Rspo1* at the onset of ovarian differentiation.

**Fig. 1.**
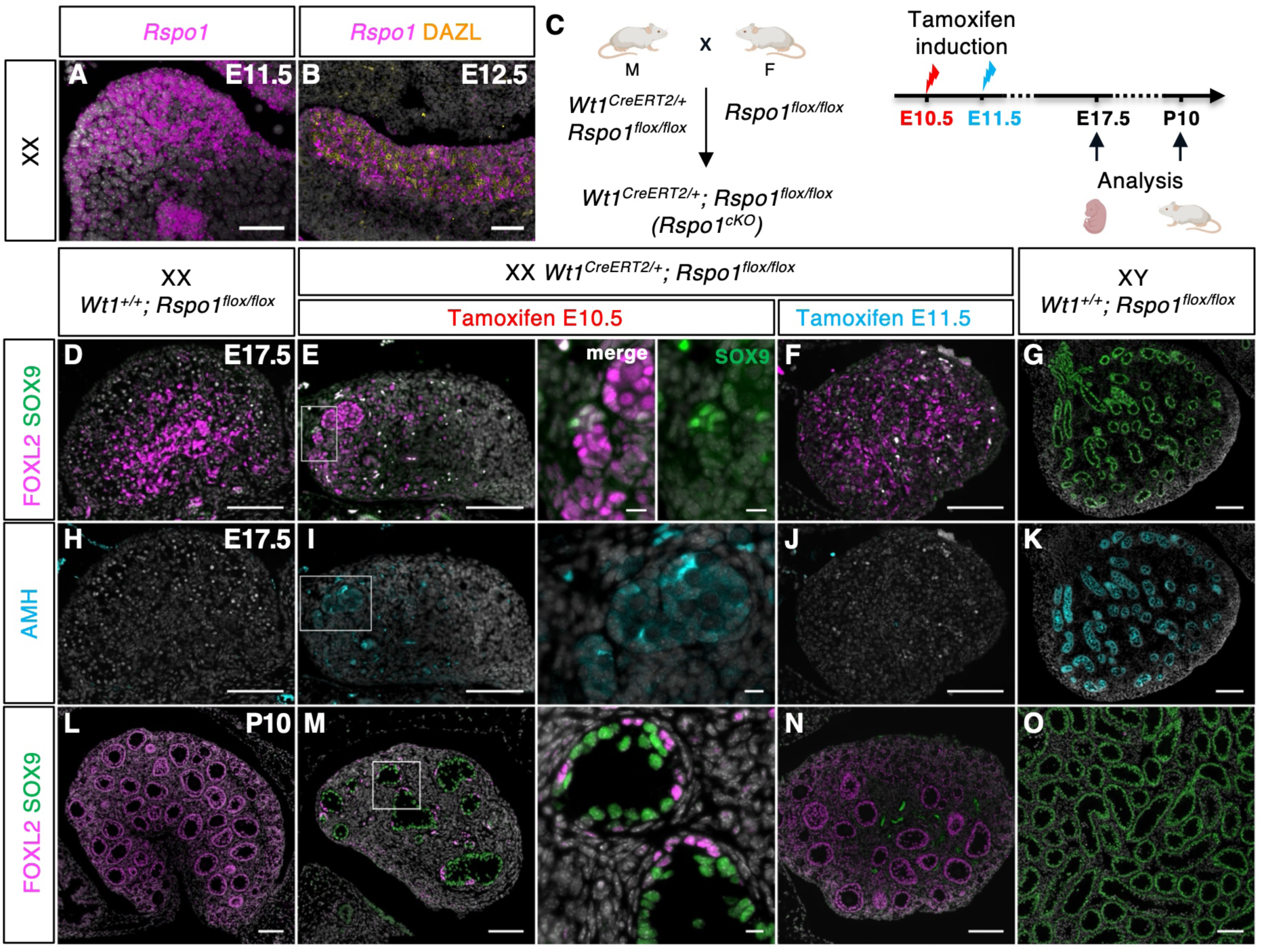
Rspo1 function is required at the onset of ovarian differentiation. **(A**) RNAscope in situ hybridization to detect *Rspo1* transcripts (magenta) in E11.5 XX gonads. (**B**) RNAscope in situ hybridization to detect *Rspo1* transcripts (magenta) and immunofluorescence for the germ cell marker DAZL (yellow) in E12.5 XX gonads. (**C**) Strategy for conditional deletion of *Rspo1* in *Wt1* positive somatic gonadal cells upon tamoxifen treatment. (**D** to **G**) Immunofluorescence for the pre-granulosa cell marker FOXL2 (magenta) and the Sertoli cell marker SOX9 (green) in the indicated genotypes at E17.5. (**H** to **K**) Immunofluorescence for the mature granulosa and Sertoli cell marker AMH (cyan) in the indicated genotypes at E17.5. (**L** to **O**) Immunofluorescence for the pre-granulosa cell marker FOXL2 (magenta) and the Sertoli cell marker SOX9 (green) in the indicated genotypes at P10. Nuclei stained with Hoechst 33342 are shown in grey. Data are representative of triplicate biological replicates. Scale bar = 100 µm. Scale bar =10 µm in zoomed insets in (E), (I) and (M).

**Fig. 2.**
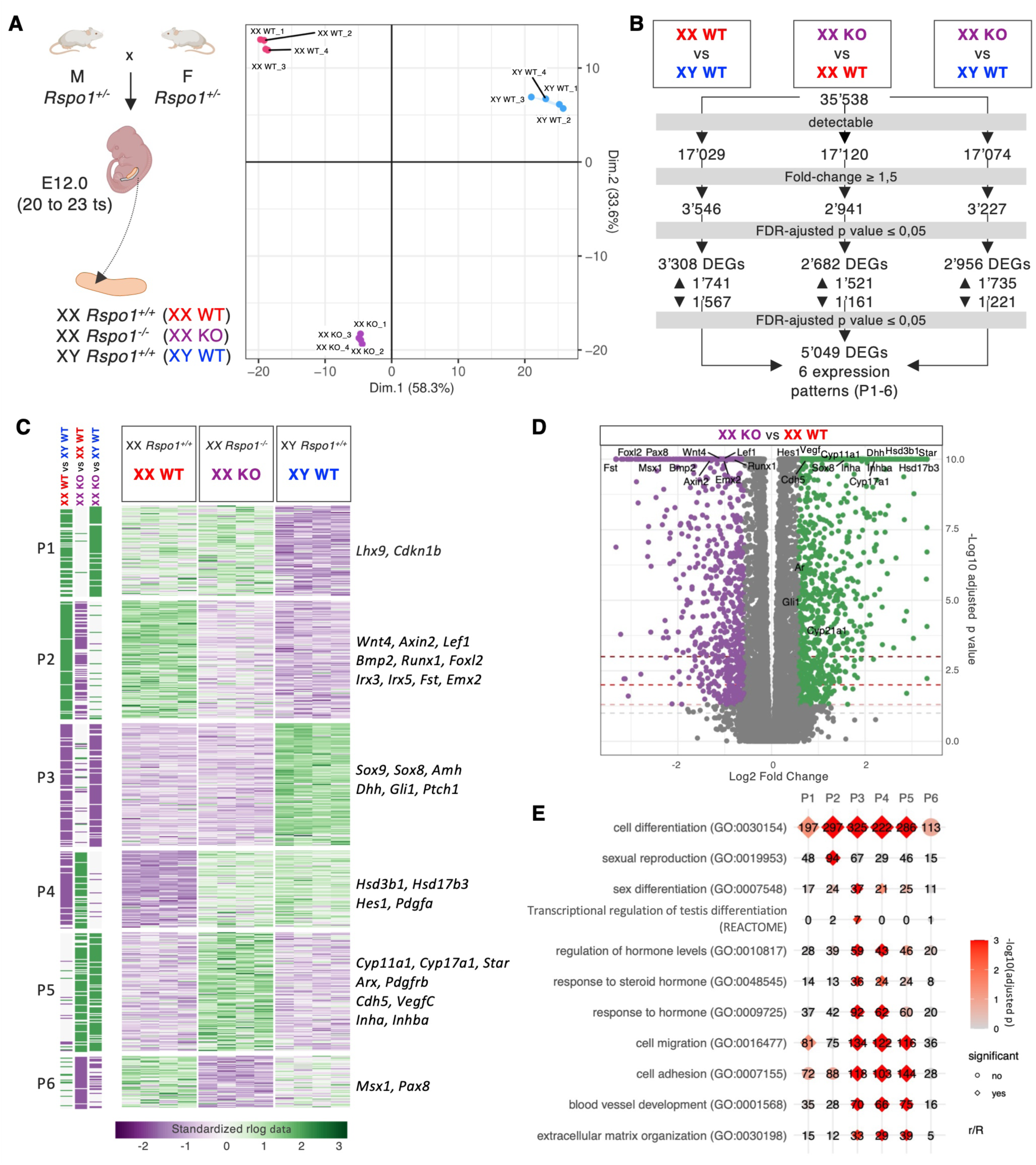
Transcriptomic analysis of *Rspo1* mutant gonads at the onset of gonadal differentiation. **(A**) Strategy and Principal Component Analysis for bulk RNA-sequencing of XX *Rspo1^+/+^* (XX WT), XX *Rspo1^-/-^* (XX KO) and XY *Rspo1^+/+^* (XY WT) E12.0 gonads. Four biological replicates (composed of 5 pairs of E12.0 gonads each) per genotype were analyzed. (**B**) Differential expression analysis pipeline. (**C**) Heatmaps representing the expression changes (standardized rlog data) of differentially expressed genes (DEGs) between XX WT, XX KO and XY WT gonads. Genes were clustered by expression profiles into 6 patterns labelled P1 to P6. Representative genes for each pattern are shown on the right side of the heatmaps. (**D**) Volcano plot shown differentially expressed genes between XX KO and XX WT gonads (-Log10 adjusted p value > 1.25). Downregulated genes (Log2 Fold Change < -0.5, purple) and up-regulated genes (Log2 Fold Change > 0.5, green) are shown. (**E**) Gene ontology enrichment analysis for DEGs in the 6 patterns.

### Transcriptomic analysis of Rspo1 mutant gonads at the onset of gonadal differentiation

To identify the initial steps leading to the masculinization in XX *Rspo1* mutant gonads, we performed bulk RNA-sequencing at E12.0. Principal Component Analysis performed using the FactoMineR package showed that the first component (PC1), accounting for 58.3% of the variance, clearly discriminated wildtype samples according to sex, whereas the second component (PC2, 33.6% of the variance) discriminated samples according to genotype, indicating that loss of *Rspo1* triggers transcriptomic changes affecting XX gonads from the onset of gonadal differentiation (Fig. 2A).

We next performed differential gene expression analysis between XX and XY wildtype samples and XX *Rspo1* mutant samples (Fig. 2B and data S1). 2682 genes were differentially expressed between XX wildtype and mutant samples highlighting the early deregulation of the ovarian transcriptional program in XX *Rspo1* mutants (Fold change ≥1.5, FDR-adjusted p value ≤ 0.05, Fig. 2, B and D, and fig. S4A).

Differentially expressed genes (DEG) partitioned into six expression patterns (Fig. 2C). Patterns 1 and 2 corresponded to genes enriched in wildtype XX gonads compared to XY gonads. While genes in pattern 1 remained detectable in XX *Rspo1* mutants, genes in pattern 2 were down-regulated, indicating that the female-associated program is impaired but not completely lost in XX *Rspo1* mutant gonads. WNT/ß-catenin targets including *Axin2*, *Lef1, Wnt4, Bmp2, Fst*, *Irx3* and *Foxl2* were down-regulated consistent with an impairment of WNT/ß-catenin signaling and pre-granulosa cell differentiation from the onset of gonadal differentiation in XX *Rspo1* mutants (Fig. 2, C and D).

Patterns 3 and 4 corresponded to genes enriched in XY gonads compared to XX gonads. Pattern 3 included genes associated to Sertoli cell differentiation such as *Sox9*, *Sox8*, *Amh* that were not detected in XX *Rspo1* mutant gonads at this stage, consistent with the known temporal dynamics of supporting cell trans-differentiation at late gestation stages (*37*). In contrast, pattern 4 captured genes associated to wildtype XY gonadal development that were up regulated in XX *Rspo1* mutant gonads at E12.0 (Fig. 2C and fig. S4A). Gene ontology analysis revealed enrichment of terms associated to cell migration (r=122, adjusted p value= 1.98E-18) and blood vessel development (66, 2.06E-14), in agreement with the formation of a male-like gonadal vasculature in XX *Rspo1* mutants (Fig. 2E and data S2) (*21*). In addition, terms associated to steroid hormone signaling (response to steroid hormone: 24, 0.006252617, regulation of hormone levels: 43, 1.64932E-06) were enriched (Fig. 2E). *Hsd3b1* and *Hsd17b3*, two genes encoding steroidogenic enzymes expressed in Fetal Leydig Cells (FLCs) and Sertoli cells respectively (*19*), were upregulated in XX *Rspo1* mutants (Fig. 2, C and D, and fig. S4A). Additional genes involved in androgen synthesis and response such as *Star, Cyp11a1*, *Cyp17a1* and *Ar*, were found in pattern 5 that comprises genes up regulated in the XX mutant gonads when compared to XX and XY controls (Fig. 2, C and D, and fig. S4A). In contrast, *Cyp19a1*, encoding the aromatase required for estrogen synthesis in mature granulosa cells, was barely detected in XX control and mutant samples at this stage (fig. S4, A and B). RT-qPCR analysis demonstrated that up-regulation of genes related to androgen synthesis in XX *Rspo1* mutant gonads was maintained at E13.5 and E17.5 (fig. S4B).

These results establish that up-regulation of gene expression related to androgen synthesis is an early and lasting signature of *Rspo1* loss of function detectable from E12.0, at least 5 days before the trans-differentiation from granulosa to Sertoli-like cell in *Rspo1* mutant ovotestes.

### Androgens production in XX Rspo1 mutant gonads

In agreement with the deregulation of steroidogenic gene expression observed in RNA-sequencing experiments, *Star* and *Cyp11a1* transcripts and HSD3B1 protein were detected in XX *Rspo1* mutant gonads at E12.5 while they were absent in control fetal ovaries (Fig. 3, B to D, and fig. S5 B to G). HSD3B1 positive cells of E16.5 XX *Rspo1* mutant gonads co-expressed *Insl3*, encoding a hormone expressed by FLCs in fetal testes and by theca cells in post-natal ovaries (*45*), highlighting their similarity to gonadal steroidogenic cells (fig. S5, K to M). In addition, E12.5 XX *Rspo1* mutant gonads expressed *Hsd17b3* transcripts, normally expressed in fetal Sertoli cells (*19*) (Fig. 3, E to G). In E16.5 XX *Rspo1* mutant gonads, *Hsd17b3* transcripts were co-expressed with *Dhh*, a marker of Sertoli cells in the fetal testis (*18*) (Fig 3I). These findings demonstrate that, in XX *Rspo1* mutant gonads gene expression related to androgen synthesis involved two distinct cell types: steroidogenic cells similar to FLCs or theca cells (expressing *Star*, *Cyp11a1*, HSD3B1 and *Insl3*) and supporting cells similar to Sertoli cells (expressing *Hsd17b3*).

**Fig. 3.**
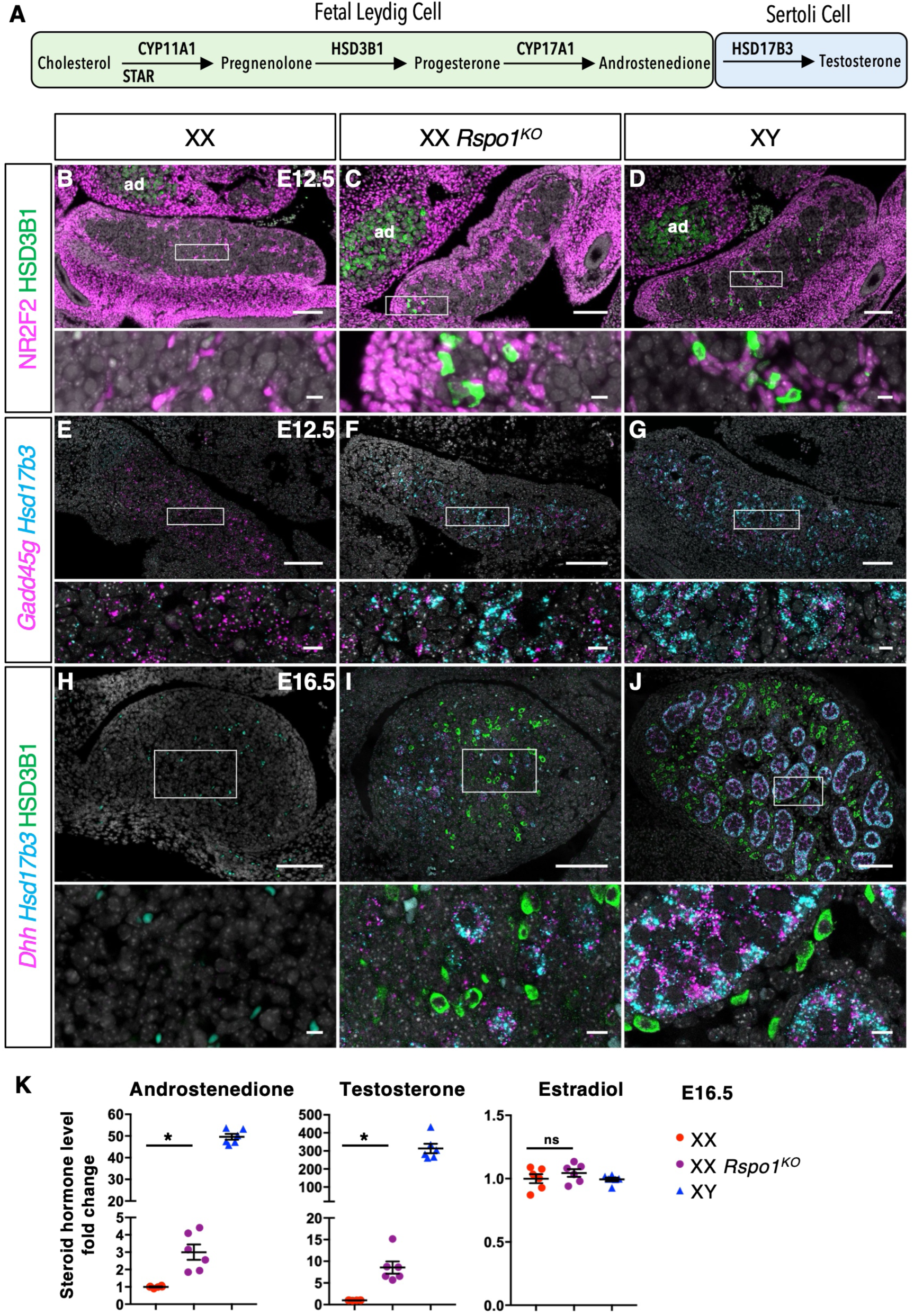
Androgens production in XX *Rspo1* mutant gonads. **(A**) Schematic representation of androgens production in the mouse fetal testis by the joined action of Fetal Leydig cells and Sertoli cells. (**B** to **D**) Immunofluorescence for the FLC marker HSD3B1 (green) and the interstitial/stromal progenitor marker NR2F2 (magenta) in the indicated genotypes at E12.5. ad: adrenal gland. (**E** to **G**) RNAscope in situ hybridization to detect transcripts for the supporting cell marker *Gadd45g* (magenta) and the Sertoli cell marker *Hsd17b3* (cyan) in the indicated genotypes at E12.5. (**H** to **J**) RNAscope in situ hybridization to detect transcripts for the Sertoli cell markers *Hsd17b3* (cyan) and *Dhh* (magenta) and immunofluorescence to detect the FLC marker HSD3B1 (green) in the indicated genotypes at E16.5. Nuclei stained with Hoechst 33342 are shown in grey. Data are representative of triplicate biological replicates. Scale bar = 100 µm. Scale bar=10 µm in zoomed insets. (**K**) Quantification of levels of androstenedione, testosterone and estradiol in E16.5 gonads determined by liquid chromatography-tandem mass spectrometry (LC-MS/MS). Fold change in steroid levels was obtained by dividing the steroid levels in gonads of a given genotype by the mean of the steroid levels in XX control gonads. N =6 pairs of E16.5 gonads per genotype. Data are shown as means ± SEM. Statistical significance was assessed by Mann-Whitney U two-tailed test. * indicates P value ≤ 0.05; ns indicates P value > 0.05.

To determine whether these cells are functional and produce androgens we performed steroid profiling by LC-MS/MS on pairs of gonads dissected from individual E16.5 embryos. Androstenedione and testosterone were detected at significant higher levels in XX *Rspo1* mutant gonads compared to XX control gonads, whereas estradiol levels were very low in all genotypes (Fig. 3K). Together these results demonstrate that abnormal steroidogenic cell differentiation from E12.5 leads to abnormal androgen synthesis in XX *Rspo1* mutant gonads.

### Androgen Receptor mediated signaling in XX Rspo1 mutant gonads

Androgens exert their biological effects by binding to the androgen receptor (AR) (*46*). *Ar* transcripts were up regulated in developing XX *Rspo1* mutant gonads (Fig. 2D and Fig. 4, F and K). Unlike control ovaries that do not express AR, XX *Rspo1* mutant gonads showed nuclear expression of AR at E14.5 (Fig. 4, A and B). AR expression was detected in NR2F2 + cells of the mutant gonads, similar to the expression found in NR2F2 + interstitial progenitors of the control testis at this stage (Fig. 4, B and C). In addition, nuclear AR was found in FOXL2+ pre-granulosa cells of the XX mutant gonads, from E14.5 (Fig. 4, D and E). At E16.5, AR was detected in the masculinized region of the mutant ovotestis in pre-granulosa cells undergoing precocious maturation marked by AMH up-regulation (Fig. 4, G to I). In contrast, AR expression was absent in pre-granulosa cells of the ovarian region of the mutant ovotestis where AMH is not expressed (Fig. 4, G to I). Known AR targets normally expressed in control mature post-natal granulosa cells (*Bmp4*, *Mmp2*) or in post-pubertal Sertoli cells (*Susd3*) were abnormally up-regulated in E17.5 XX *Rspo1* mutant gonads (Fig. 4K) (*47, 48*). These results indicate that androgens produced in XX *Rspo1* mutant gonads can signal through AR to trigger its nuclear translocation and activate downstream signaling.

**Fig. 4.**
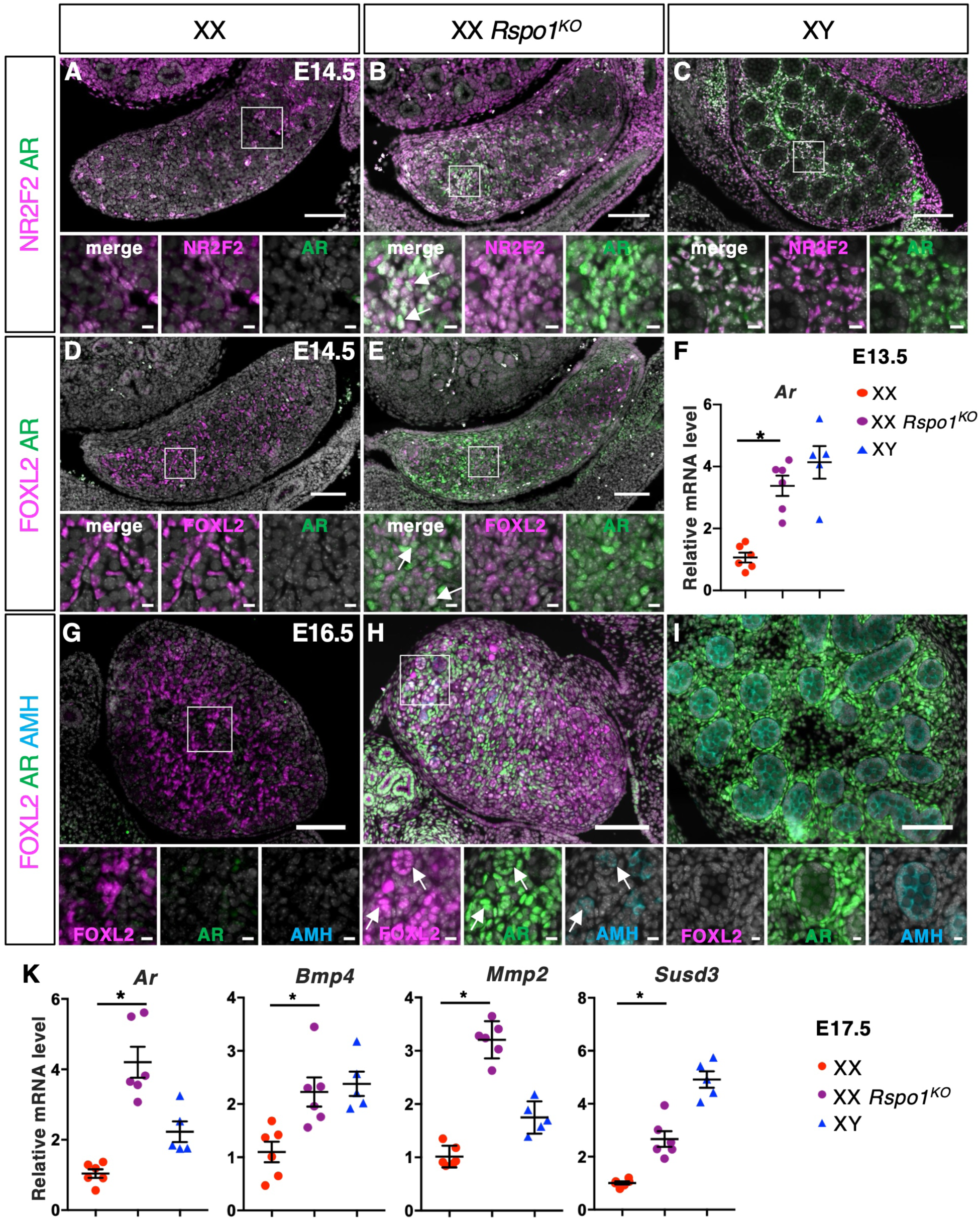
Androgen Receptor mediated signaling in XX *Rspo1* mutant gonads. (**A** to **C**) Immunofluorescence for the interstitial/stromal progenitor marker NR2F2 (magenta) and Androgen Receptor (AR, green) in the indicated genotypes at E14.5. Arrows in insets indicate double positive cells. (**D** to **E**) Immunofluorescence for the pre-granulosa cell marker FOXL2 (magenta) and AR (green) in the indicated genotypes at E14.5. Arrows in merge inset in (**E**) indicate double positive cells. (**F**) Quantification of *Ar* transcripts at E13.5 after normalization to *Sdha* and *Tbp* by RT-qPCR. (**G** to **I**) Immunofluorescence for the pre-granulosa cell marker FOXL2 (magenta), AR (green) and the mature granulosa cell and Sertoli cell marker AMH (cyan) in the indicated genotypes at E16.5. Arrows in insets in (**H**) indicate triple positive cells. (**K**) Quantification of *Ar*, *Bmp4, Mmp2* and *Susd3* transcripts at E17.5 after normalization to *Sdha* and *Tbp* by RT-qPCR. Immunofluorescence data are representative of triplicate biological replicates. Nuclei stained with Hoechst 33342 are shown in grey. Scale bar = 100 µm. Scale bar=10 µm in zoomed insets. RT-qPCR data are shown as means ± SEM. Statistical significance was assessed by Mann-Whitney U two-tailed test. * indicates P value ≤ 0.05; ns indicates P value > 0.05.

### Inhibition of Androgen signaling rescues ovarian development in XX Rspo1 mutant gonads

To decipher the role of androgen signaling in the masculinization phenotype of XX *Rspo1* mutant gonads, we inhibited pharmacologically AR function by treating pregnant females with the nonsteroidal AR antagonist flutamide (Fig. 5A) (*25*). Ano-genital distance in XY embryos, a readout of androgen signaling, was reduced following flutamide treatment, consistent with decreased AR signaling (fig. S6B). In addition, while vehicle-treated XX *Rspo1* mutants exhibited pseudo-hermaphroditism of the internal genitalia, flutamide-treated XX *Rspo1* mutants developed female internal genitalia without male-like structures (fig. S6, D and F) (*25*). Together, these findings demonstrate that maternal flutamide treatment effectively reduces AR signaling in developing embryos.

**Fig. 5.**
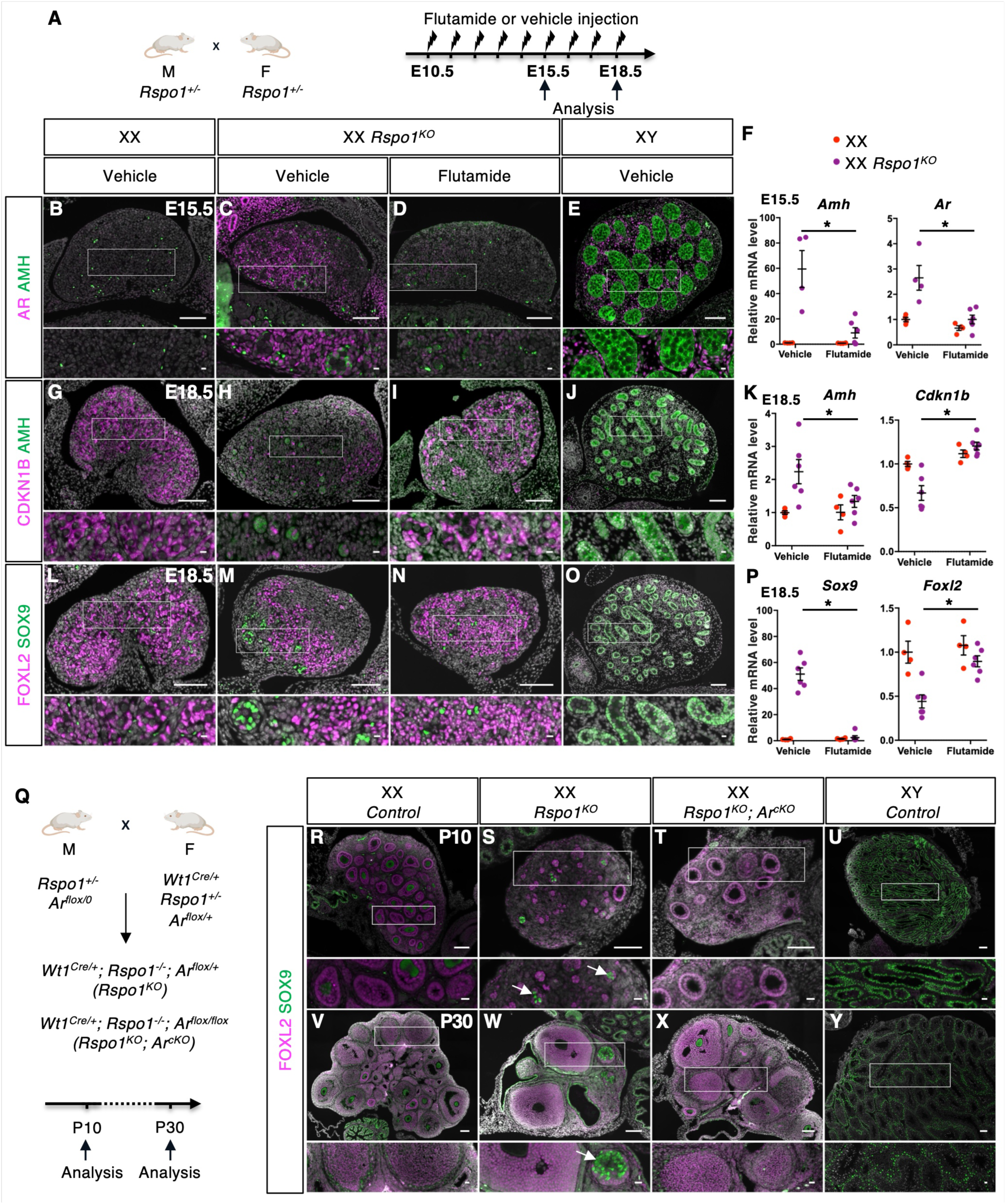
Inhibition of Androgen signaling rescues ovarian development in XX *Rspo1* mutant gonads. (**A**) Strategy for flutamide or vehicle treatment of *Rspo1^+/-^* pregnant females. (**B** to **E**) Immunofluorescence for AR (magenta) and the mature granulosa cell and Sertoli cell marker AMH (green) in the indicated genotypes at E15.5 (**F**) Quantification of *Amh* and *Ar* transcripts at E15.5 after normalization to *Sdha* and *Tbp* by RT-qPCR. (**G** to **J**) Immunofluorescence for the mitotically arrested pre-granulosa cell marker CDKN1B (magenta) and the mature granulosa cell and Sertoli cell marker AMH (green) in the indicated genotypes at E18.5. (**K**) Quantification of *Amh* and *Cdkn1b* transcripts at E18.5 after normalization to *Sdha* and *Tbp* by RT-qPCR. (**L** to **O**) Immunofluorescence for the pre-granulosa cell marker FOXL2 (magenta) and the Sertoli cell marker SOX9 (green) in the indicated genotypes at E18.5. (**P**) Quantification of *Sox9* and *Foxl2* transcripts at E18.5 after normalization to *Sdha* and *Tbp* by RT-qPCR. (**Q**) Strategy to generate *Rspo1^KO^; Ar^cKO^*double mutants. (**R** to **U**) Immunofluorescence for FOXL2 (magenta) and SOX9 (green) in the indicated genotypes at P10. White arrows in the zoomed areas in (**S**) indicate SOX9+ cells. (**V** to **Y**) Immunofluorescence for FOXL2 (magenta) and SOX9 (green) in the indicated genotypes at P30. White arrow in the zoomed area in (**W**) indicates seminiferous tubule-like structure. Immunofluorescence data are representative of triplicate biological replicates. Nuclei stained with Hoechst 33342 are shown in grey. Scale bar = 100 µm. Scale bar=10 µm in zoomed insets. RT-qPCR data are shown as means ± SEM. Statistical significance was assessed by Mann-Whitney U two-tailed test. * indicates P value ≤ 0.05; ns indicates P value > 0.05.

Next, we examined the impact of flutamide treatment on gonadal development in XX *Rspo1* mutants (Fig. 5, A to P). Vehicle-treated mutant gonads developed as ovotestes with precocious maturation of granulosa cells evidenced by the up-regulation of *Amh*/AMH and *Ar*/AR at E15.5 followed by the down-regulation of the mitotically arrested pre-granulosa cell marker *Cdkn1b*/CDKN1B (*37*) (Fig. 5, C, F, H and K). At E18.5, granulosa cells were undergoing trans-differentiation into Sertoli-like cells, as indicated by the down-regulation of *Foxl2*/FOXL2 and the up-regulation of *Sox9*/SOX9 (Fig. 5, M and P). In contrast, flutamide-treated XX *Rspo1* mutant gonads showed reduced expression of *Amh*/AMH and *Ar*/AR from E15.5 (Fig. 5, D and F) as well as increased expression of *Cdkn1b*/CDKN1B and *Foxl2*/FOXL2 and reduced *Sox9*/SOX9 expression at E18.5 (Fig. 5, I, K, N and P). Together these observations indicate that premature differentiation of granulosa cells and their subsequent trans-differentiation into Sertoli-like cells in XX *Rspo1* mutant gonads are dependent on androgen signaling and can be prevented by pharmacological inhibition of AR.

To completely abolish androgen receptor function in the context of the *Rspo1* mutation we adopted a genetic approach and generated XX *Rspo1^KO^*; *Ar^cKO^* double mutants (Fig. 5Q). Efficient deletion of a conditional *Ar* allele in *Wt1* expressing tissues, including gonadal somatic cells, was achieved using the *Wt1^Cre^*line (*42, 49*) (fig. S6, K to N). We compared the phenotypes of XX *Rspo1^KO^*single mutants (*Wt1^Cre/+^; Rspo1^-/-^; Ar^flox/+^*) and XX *Rspo1^KO^*; *Ar^cKO^* double mutants (*Wt1^Cre/+^; Rspo1^-/-^; Ar^flox/flox^*) at P10 and P30. XX *Rspo1^KO^*single mutants exhibited pseudo-hermaphroditism of the internal genitalia (fig. S6, G and H) and gonads that developed as ovotestes as shown by the presence of SOX9+ Sertoli-like cells organized into seminiferous tubule-like structures (Fig. 5, S and W). XX *Rspo1^KO^*; *Ar^cKO^* double mutants developed female internal genitalia without male-like structures (fig. S6I). In contrast to the single mutants, XX *Rspo1^KO^*; *Ar^cKO^*double mutant gonads lacked SOX9+ Sertoli-like cells and seminiferous tubule-like structures and harbored abundant follicles (Fig. 5, T and X). These findings demonstrate that ablation of the androgen receptor rescues the ovotestis phenotype of XX *Rspo1* mutants indicating that deregulation of intra-gonadal androgen production and signaling are major drivers of gonadal masculinization in XX *Rspo1* mutants.

## Discussion

Mouse ovarian development requires high levels of WNT/ß-catenin signaling mediated by RSPO1 as mutations in *Rspo1*, *Wnt4* and *Ctnnb1* lead to XX sex reversal with ovotesticular development (*21, 22, 24, 25*). We have established that RSPO1 function is required in somatic gonadal cells before E12.5, demonstrating that timely RSPO1/WNT/ß-catenin signaling is crucial for ovarian development. Our findings reveal abnormal steroidogenic gene expression and androgen production in *Rspo1* mutants consistent with previous observations in XX gonads mutant for *Wnt4* and *Ctnnb1* (*22, 24, 50*), highlighting that RSPO1/WNT/ß-catenin signaling plays an essential role in suppressing androgen production in developing ovaries.

Notably, inhibition of androgen receptor function rescues ovarian sex reversal, establishing androgen signaling as a key driver of ovarian masculinization in *Rspo1* mutants, in addition to its role in genitalia virilization. Although androgen signaling does not affect the development of the wildtype fetal ovary (*7–9*), disruption of RSPO1/WNT/β-catenin signaling enables ovarian somatic cells to produce and respond to androgens, thereby promoting ovarian masculinization. These findings demonstrate that RSPO1/WNT/β-catenin signaling protects the mouse fetal ovary against androgen-dependent sex reversal by inhibiting both the production and the ability to respond to androgens.

This concept may also be relevant to human ovarian development. In humans, folliculogenesis is initiated in utero from 13–14 PCW (*51*). Androgens levels in human fetal ovaries are very low, whereas the estrogen levels are relatively high (*31, 32*). This suggests that most of the androgens produced by theca cells are converted into estrogens by CYP19A1, which is expressed in the granulosa cells of developing follicles (*52*). Consistent with findings from genetic mouse models, mutations in *RSPO1* and *WNT4* can lead to 46,XX testicular or ovotesticular DSD (*33, 34, 53*). The masculinization of the external genitalia in these patients indicates increased androgen production during fetal life. As *RSPO1* deficiency is not lethal, the gonadal phenotype has primarily been characterized in adult 46,XX patients, in whom Leydig cell hyperplasia has been reported (*33*). In contrast, *WNT4* mutations can be lethal, and Leydig cells have been described in the gonads of XX fetus with a *WNT4* mutation (*53*). These observations support a conserved role for RSPO1/WNT/ß-catenin pathway in suppressing androgen production and androgen-dependent masculinization in the human fetal ovary. Thus, androgen resistance is actively maintained by RSPO1/WNT/β-catenin signaling as part of the mammalian ovarian developmental program.

## Acknowledgments

We thank the Institut Clinique de la Souris – PHENOMIN-ICS for their expert assistance in establishing the conditional *Rspo1* mutant mouse line and Pr. Franck Claessens for sharing the *Ar^tm1Verh^* line. We acknowledge the help from members of the Experimental Histopathology Platform, the PRISM Imaging Platform and the Animal house at iBV (Institut de Biologie Valrose, Université Côte d’Azur, CNRS, Inserm, iBV, France). We are grateful to members of the A. Schedl, and M.C. Chaboissier groups for helpful discussions.

## Funding

Agence Nationale de la Recherche ANR-19-CE14-0022 SexDiff (MCC, FCh)

Agence Nationale de la Recherche ANR-23-CE14-0012, Heterosex (MCC)

Fondation Maladies Rares, Phenomin 2013 Mouse models and rare diseases (MCC)

China Scholarship Council 201506300094 (FT)

Deutsche Forschungsgemeinschaft (DFG) Instrument grant support INST 269/910-1 FUGG (MP)

## Author contributions

Conceptualization: FT, FCh, MCC, APG

Investigation: FT, NBC, KB, AL, TAD, MD, AD, MP, MCB, FCh, APG

Visualization: FT, FCh, APG

Funding acquisition: MP, FCh, MCC

Supervision: MCC, APG

Writing – original draft: FT, FCh, MCC, APG

Writing – review & editing: FT, NBC, KB, MD, MP, MCB, FCh, MCC, APG

## Data, code, and materials availability

All data are available in the main text or the supplementary materials.

The RNA-sequencing data discussed in this publication have been deposited in NCBI’s Gene Expression Omnibus and are accessible through GEO Series accession number GSEXXX.

Functional enrichment analysis used an R implementation derived from the AMEN suite of tools) publicly available at https://sourceforge.net/projects/amen/.

Materials generated in this study are available from the corresponding author upon reasonable request. The *Ar^tm1Verh^* line was obtained from KU Leuven under a Material Transfer Agreement (MTA) and cannot be redistributed by the authors. Requests for access to this line should be directed to KU Leuven.

### Supplementary Materials

Materials and Methods

Figs. S1 to S6

Tables S1 to S3

References cited in Supplementary Materials: 54 to 72.

**Other Supplementary Materials for this manuscript include the following:**

Data S1 and S2

## Materials and Methods

### Establishment of a new *Rspo1* exon 2 conditional deletion mutant line

The *Rspo1* knock-out first (with conditional potential) mutant mouse line was generated at the *Institut Clinique de la Souris – PHENOMIN* (http://www.phenomin.fr). An embryonic stem (ES) cell clone, EPD0720_2_D03, derived from the parental JM8A3.N1 line and produced within the framework of the KOMP program, was obtained from the Mutant Mouse Resource and Research Center (MMRRC). This clone was validated by Southern blotting using an internal *Neo* probe and by PCR confirming the presence of the 3′ *LoxP* site. Following verification of a normal karyotype by chromosome spreading and Giemsa staining, the clone was microinjected into BALB/cN blastocysts. Male chimeras were obtained, and germline transmission was successfully achieved for the *Rspo1^tm2a(KOMP)Wtsi^*allele (knock-out first with conditional potential). The conditional *Rspo1^tm2c(KOMP)Wtsi^*allele (referred to as *Rspo1^flox^*) where *Rspo1* exon 2 is flanked by LoxP sites, was obtained by breeding *Rspo1^tm2a(KOMP)Wtsi^*heterozygous animals with the *Tg(CAG-flpo)1Afst* Flp-deleter line (*54*).

### Establishment of a new *Rspo1* exon 2 constitutive deletion mutant line

In order to obtain a germ line deletion of *Rspo1* exon 2 we used the *Edil3^Tg(Sox2-cre)1Amc^* transgenic line where Cre is expressed in oocytes under the control of Sox2 regulatory sequences (referred to as *Sox2:Cre^tg^*) (*55, 56*). *Rspo1^flox/flox^*males were crossed with *Sox2:Cre^tg/0^*females. We recovered *Rspo1^+/-^* individuals where *Rspo1* exon 2 had been deleted in zygotes by the maternally expressed Cre.

### Additional mouse strains

The experiments described herein were carried out in compliance with the guidelines of the French Regulations for Animal Care and with the approval of the local Ethical Committee (APAFIS APAFIS#12789-2017121515109323v1 and APAFIS#15904-2018061816265103v7). Mouse lines were kept on a mixed background B6CBAF1/JRj. The knock-in *Wt1^tm2(cre/ERT2)Wtp^* line (referred to as *Wt1^CreERT2^*) (*42*) was used to generate *Rspo1* conditional mutants (*Rspo1^cKO^*). *Wt1^CreERT2/+^; Rspo1^flox/flox^* males were crossed with *Rspo1^flox/flox^* females to obtain *Wt1^CreERT2/+^; Rspo1^flox/flox^* embryos (*Rspo1^cKO^*) or *Rspo1^flox/flox^*embryos (controls). To activate the Cre^ERT2^ recombinase in embryos, tamoxifen (T5648, Sigma-Aldrich) was directly diluted in corn oil to a concentration of 40 mg/mL, and tamoxifen treatments (200 mg/kg body weight) were administered to pregnant females by oral gavage at E10.5 or E11.5. Tamoxifen treatment causes dystocia in pregnant dams; therefore, to obtain P10 samples, pregnant dams were delivered by cesarean section at E18.5, and the newborn pups were fostered and nursed by surrogate mothers. The transgenic *Tg(Nr5a1-cre)2Klp* line (referred to as *Nr5a1-Cre*) (*43*) was used to delete *Rspo1* in somatic gonadal cells from E12.5 (*44*). The knock-in *Wt1^tm1(EGFP/cre)Wtp^* line (referred to as *Wt1^Cre^*) (*42*), was crossed with the *Ar^tm1Verh^* line (referred to as *Ar^flox^*) (*49*) for conditional deletion of *Ar* (*Ar^cKO^*). *Rspo1^+/-^; Ar^flox/0^* males were crossed with *Wt1^Cre/+^; Rspo1^+/-^; Ar^flox/+^* females to generate XX *Rspo1^KO^* mutants (*Wt1^Cre/+^; Rspo1^-/-^; Ar^flox/+^*), XX *Rspo1^KO^*; *Ar^cKO^* double mutants (*Wt1^Cre/+^; Rspo1^-/-^; Ar^flox/flox^*) and XX controls (*Wt1^Cre/+^; Rspo1^+/-^; Ar^flox/+^* or *Wt1^Cre/+^; Rspo1^+/+^; Ar^flox/+^*). The day when a vaginal plug was found was designated as embryonic day E0.5. Embryos dissected on days 11 and 12 of gestation were staged by counting the number of tail somites (ts).

### Genotyping

Genotypes of mice and embryos were determined using PCR assays on lysates from ear biopsies or tail tips. Genotyping of the *Rspo1* exon 2 conditional and mutant alleles was achieved by PCR using *Rspo1-Ef* and *Rspo1-Er* primers (table S1) to amplify a 300 bp band for the wildtype *Rspo1* allele and a 450 bp band for the *Rspo1^flox^* allele, and the *Rspo1-Ef* and *Rspo1-Lxr* primers (table S1) to amplify a 300 bp band for the *Rspo1^-^*allele. Genotyping of Flp deleter line, *Sox2:Cre^tg^*, *Wt1^CreERT2^*, *Wt1^Cre^*, *Nr5a1-Cre* and *Ar^flox^* lines was performed as described in (*42, 43, 49, 54, 55*). Genotyping primers are listed in Table S1.

### Flutamide treatment and anogenital distance measurements

Flutamide (F9397, Sigma-Aldrich) was prepared in corn oil containing 15% ethanol to concentration of 100mg/mL at 60°C. Pregnant *Rspo1^+/-^*mice were randomly distributed in two groups and were injected subcutaneously at the dose of 200 mg/kg body weight of flutamide or the equivalent volume of vehicle (15% ethanol in oil) once daily from E10.5 to E15.5 or from E10.5 to E18.5. Anogenital distance (distance between the anus and the external genitalia) was measured in E18.5 fetuses with an electronic caliper under the dissecting microscope. Weight and anogenital distance measurements of embryos after flutamide/vehicle treatments were performed blindly before genotyping.

### Immunofluorescence and RNAscope situ hybridization

Embryos were fixed in 4% (*w/v*) paraformaldehyde (PFA, 15710-S, EMS) overnight, processed for paraffin embedding, and sectioned into 5 µm thick sections. Immunofluorescence and Hoechst 33342 (H3570, Invitrogen) staining were performed as described in (*57*). mRNA was detected with the RNAscope^TM^ technology following manufacturer’s instructions using the RNAscopeMultiplex Fluorescent Reagent Kit v2 Assay (Advanced Cell Diagnostics). Epifluorescence images were obtained on a motorized Axio Imager Z1 microscope (Zeiss) coupled with an AxioCam MRm camera (Zeiss). Confocal images were obtained on a Zeiss LSM 780 microscope using a 40x/1.2 W. Images were processed with Fiji (Bethesda, MD, USA) and assembled using the open-source software platform OMERO (https://www.openmicroscopy.org/omero/). Antibodies and RNAscope probes are listed in Table S2. At least three embryos of each genotype were analyzed for each marker.

### RNA extraction and quantitative PCR analyses

Individual gonads were dissected from the mesonephros in PBS, snap-frozen in liquid nitrogen and kept at −80 °C. RNA extraction, cDNA preparation and quantitative PCR were performed as described in (*58*). Primer sequences are listed in Table S1. All biological replicates of different genotypes (*N* = 4–6) were run in the same plate and run as duplicate technical replicates. Relative gene expression of each gonad was normalized to the expression of the housekeeping genes *Shda* and *Tbp* (*59*) by the 2^-ΔΔCT^ calculation method. GeNorm, BestKeeper algorithms and the comparative delta-CT method provided through the online tool RefFinder (https://www.ciidirsinaloa.com.mx/RefFinder-master/?type=reference#) were used to confirm reference gene stability in the experimental datasets. Fold change in gene expression was obtained by dividing the normalized gene expression in gonads of a given genotype by the mean of the normalized gene expression in XX control gonads. Data are shown as means ± SEM. Statistical significance was assessed by Mann-Whitney U two-tailed test (GraphPad Prism 10.2.1). * indicates P value ≤ 0.05; ns indicates P value > 0.05.

### Generation of cDNA libraries for bulk RNA-sequencing of *Rspo1* mutant gonads

Pairs of gonads from E12.0 (20 to 23 tail somites) individual embryos were dissected from the mesonephros in PBS, snap-frozen in liquid nitrogen and kept at −80 °C. 5 pairs of gonads of identical genotype were pooled and RNA was extracted using RNeasy Micro Kit (74004, Qiagen). 4 replicates containing 5 pairs of gonads were used for XX *Rspo1^+/+^*, XX *Rspo1^-/-^* and XY *Rspo1^+/+^* samples. Library preparation and sequencing were performed by Novogen. Briefly, RNA integrity and quantitation were assessed using RNA 6000 Nano Kit on the Bioanalyzer 2100 system (Agilent Technologies). A total amount of 1 μg RNA per sample was used as input material for the RNA sample preparations. Sequencing libraries were generated using NEBNext Ultra RNA Library Prep Kit for Illumina (NEB, USA) following manufacturer’s recommendations and index codes were added to attribute sequences to each sample. Briefly, mRNA was purified from total RNA using poly-T oligo-attached magnetic beads. Fragmentation was carried out using divalent cations under elevated temperature in NEBNext First Strand Synthesis Reaction Buffer (5X). First strand cDNA was synthesized using random hexamer primer and M-MuLV Reverse Transcriptase (RNase H-). Second strand cDNA synthesis was subsequently performed using DNA Polymerase I and RNase H. Remaining overhangs were converted into blunt ends via exonuclease/polymerase activities. After adenylation of 3’ ends of DNA fragments, NEBNext Adaptor with hairpin loop structure were ligated to prepare for hybridization. To select cDNA fragments of preferentially 150∼200 bp in length, the library fragments were purified with AMPure XP system (Beckman Coulter, Beverly, USA). Then 3 μl USER Enzyme (NEB, USA) was used with size-selected, adaptor-ligated cDNA at 37 °C for 15 min followed by 5 min at 95 °C before PCR. Then PCR was performed with Phusion High-Fidelity DNA polymerase, Universal PCR primers and Index (X) Primer. At last, PCR products were purified (AMPure XP system) and library quality was assessed on the Agilent Bioanalyzer 2100 system. The clustering of the index-coded samples was performed on a cBot Cluster Generation System using PE Cluster Kit cBot-HS (Illumina) according to the manufacturer’s instructions. After cluster generation, the library preparations were sequenced on an Illumina platform and paired-end reads were generated.

### Analysis of bulk RNA-sequencing of *Rspo1* mutant gonads

#### Data preprocessing

Paired-end reads were adapter- and quality-trimmed with TrimGalore v0.6.11 (Cutadapt v5.2; --paired, -q 20, minimum length 20 bp) (*60, 61*) and then, quantified with Salmon v1.11.4 (*62*) in selective-alignment mode against a decoy-aware index of the Ensembl release 115 mouse transcriptome (GRCm39), with libraries treated as unstranded (--libType A). Finally, per-sample read counts were merged into a single count matrix (outer join, missing values set to 0).

#### Detecting and filtering out low-quality samples

To identify and exclude low-quality samples, those with a total read count below 10% of the overall median were flagged for exclusion. This approach accounts for inherent variability across different experiments and sequencing platforms, ensuring more reliable and consistent downstream analyses. No samples were excluded from our study.

#### Detecting and filtering lowly expressed genes

Genes were pre-filtered using the filterByExpr function from edgeR (v. 4.6.2; (*63*)) with min.count = 3 and min.total.count = 10; genes with zero variance were excluded.

#### Data normalization

To stabilize variance and normalize count distributions across samples, the regularized log transformation (rlog) from the DESeq2 package (*64*) was applied. All RNA-seq data, both raw and preprocessed are available at the GEO repository (*65*) under the accession number GSEXXXX.

#### Differential gene expression analysis

RNA-seq differential gene expression analysis was performed as described by (*66*). Briefly, four statistical methods were applied: limma-voom (v. 3.64.1; (*67*), edgeR (v. 4.6.2; (*63*), DESeq2 (v. 1.48.1; (*64*) and RankProducts (v. 3.34.0; (*68*). Within each comparison, p-values from the four tools were integrated using Stouffer’s method (also called Stouffer-Lipták) (*69*). All p-values were adjusted for multiple testing by the Benjamini–Hochberg procedure (*70*). Genes were considered significantly differentially expressed in a given comparison if the adjusted p-value was below 0.05 and the absolute log2 fold-change exceeded log2(1.5). To increase the statistical power and robustness of the identified genes across multiple comparisons, the resulting Stouffer p-values were further combined using Fisher’s method. Genes were selected as differentially expressed across comparisons if the adjusted p-value was below 0.05 and the absolute fold-change exceeded 1.5 in at least one individual comparison.

#### Clustering analysis

The resulting differentially expressed genes (DEGs) were partitioned into expression patterns using the k-means algorithm (*71*). Expression profiles were displayed using the R package pheatmap (Kolde, 2010, pheatmap: Pretty Heatmaps. R package version 1.0.13, https://cran.r-project.org/web/packages/pheatmap).

#### Functional enrichment analysis

Functional enrichment analyses were performed using an R implementation derived from the AMEN suite of tools (*72*). Enrichment p-values were computed using the hypergeometric distribution. A term was retained if the Benjamini–Hochberg adjusted p-value was ≤0.05 (*70*) and at least three DEGs mapped to it.

#### LC-MS/MS analysis of steroid hormones

Pairs of gonads from E16.5 individual embryos were dissected from the mesonephros in PBS, snap-frozen in liquid nitrogen and kept at −80 °C. Hormones levels of androstenedione, testosterone and estradiol were determined by liquid chromatography-tandem mass spectrometry (LC-MS/MS) using a QTRAP 6550+ (Sciex, Darmstadt) coupled to an Aquitiy i-class ultra-performance liquid chromatography system (Waters, Eschborn). Samples were extracted in cold methanol including respective stable isotope labelled internal standards for 10min using a cooled ultrasonic bath. After homogenization sample extracts were dried down under a gentle stream of nitrogen. For analysis, samples were reconstituted in a mixture of 75%/25% (v/v) mobile phases A (0.2mM ammonium fluoride) and B (methanol). Chromatographic separation was achieved at 40°C using a Phenomenex Kinetex C18 EVO column (150x2.1mm, 2.6µm; Phenomenex, Aschaffenburg). Therefore, samples were injected into a flow rate of 0.35mL/min of 75% mobile phase A, followed by a linear decrease to 51% at 3.0min and further down to 0% at 8.0min. After a hold for 1min, initial conditions were reset within 0.1min followed by a re-equilibration of the LC system. Androstenedione and testosterone were analyzed by positive electrospray ionization (ESI; ion source voltage 4500V) using the multiple reaction scan mode (MRM), whereas estradiol was analyzed in negative ESI (ion source voltage -4500V). As precursor/Q1 masses, 287.1, 289.1 and 271.2 were used for androstenedione, testosterone and estradiol, respectively. As product/Q3 ions 109.1 and 97.1 were used for androstenedione and testosterone, whereas 145.1 and 183.3 were used for detection of estradiol. Quantification of steroid levels was done by comparisons of ratios of analyte peak area obtained from samples to respective peak areas of stable isotope labelled internal standards to those observed in calibrators. Fold change in steroid levels was obtained by dividing the steroid levels in gonads of a given genotype by the mean of the steroid levels in XX control gonads. 6 pairs of gonads from 6 individuals per genotype (XX *Rspo1^+/+^*, XX *Rspo1^-/-^* and XY *Rspo1^+/+^*) were analyzed. Data are shown as means ± SEM. Statistical significance was assessed by Mann-Whitney U two-tailed test (GraphPad Prism 10.2.1). * indicates P value ≤ 0.05; ns indicates P value > 0.05.

---

**Fig. S1.**
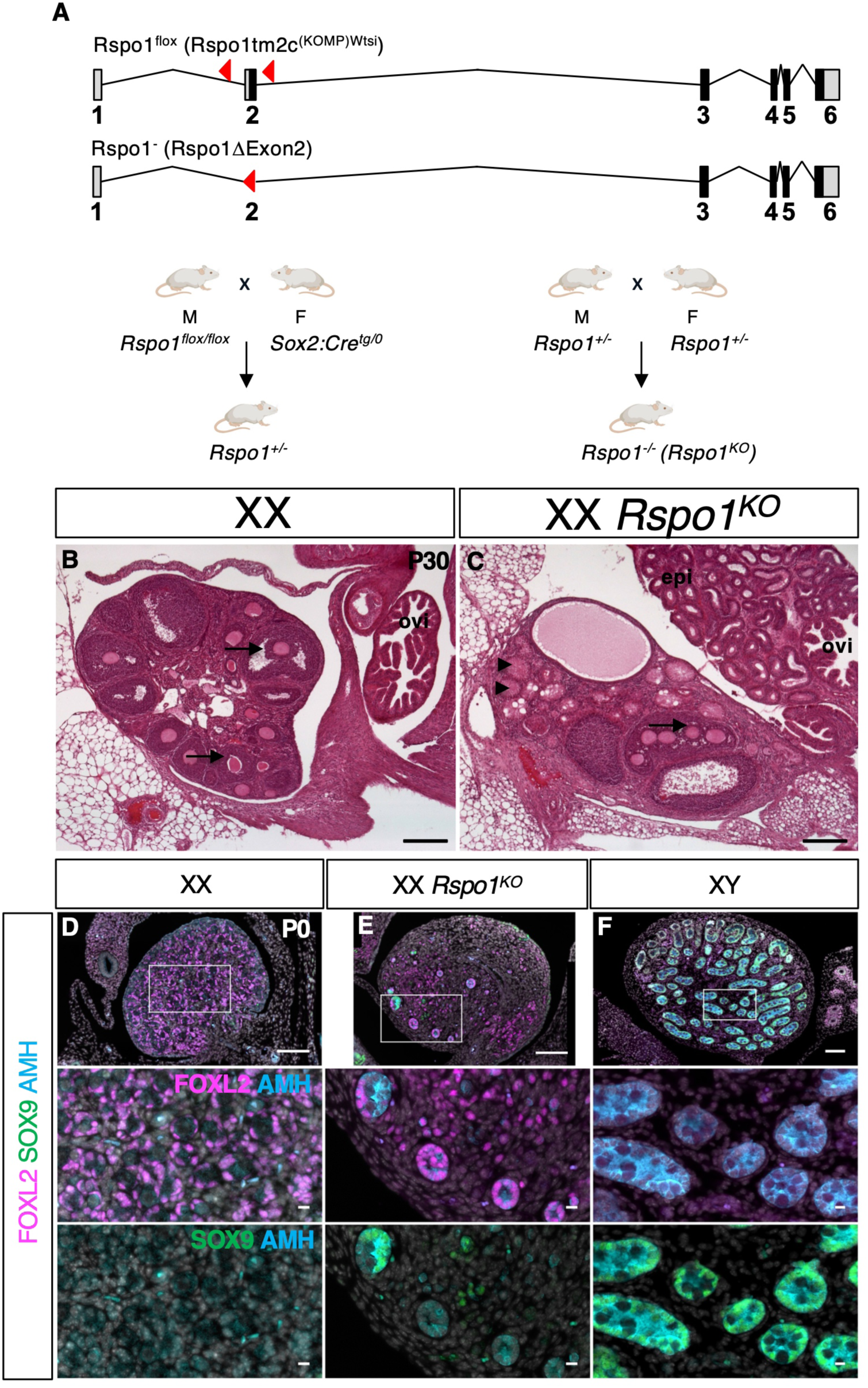
Validation of a new conditional mutant allele for deletion of *Rspo1* exon 2. (B) In *Rspo1^flox^* allele (*Rspo1tm2c^(KOMP)Wtsi^*), exon 2 of *Rspo1* is flanked by two LoxP sites. Upon Cre mediated recombination, exon 2 (containing the ATG and the peptide signal) is excised resulting in a loss of function allele (*Rspo1^-^*). *Rspo1^flox/flox^* males were crossed with *Sox2:Cre ^tg/0^* females to generate a germ line deletion of *Rspo1* exon 2. The resulting *Rspo1^+/-^*individuals were inter-crossed to generate *Rspo1^-/-^* (*Rspo1^KO^*) individuals. (**B** and **C**) Hematoxylin/eosin staining of sections of XX Wildtype and XX *Rspo1^KO^* gonads at post-natal day 30 (P30). XX Wildtype gonad is an ovary containing developing follicles (arrows). XX *Rspo1^KO^* gonad is an ovotestis containing both follicles (arrows) and seminiferous tubule-like structures (arrowheads). XX *Rspo1^KO^* individuals exhibit internal genitalia pseudo-hermaphroditism with the development of both male structures (epi: epididymis) and female structures (ovi: oviduct). Data are representative of triplicate biological replicates. Scale bar = 200 µm. (**D** to **F**) Immunofluorescence for the pre-granulosa cell marker FOXL2 (magenta), the Sertoli cell marker SOX9 (green) and the mature granulosa and Sertoli cell marker AMH (cyan) in the indicated genotypes at P0. Nuclei stained with Hoechst 33342 are shown in grey. Data are represntative of triplicate biological replicates. Scale bar = 100 µm. Scale bar = 10 µm in zoomed insets.

**Fig. S2.**
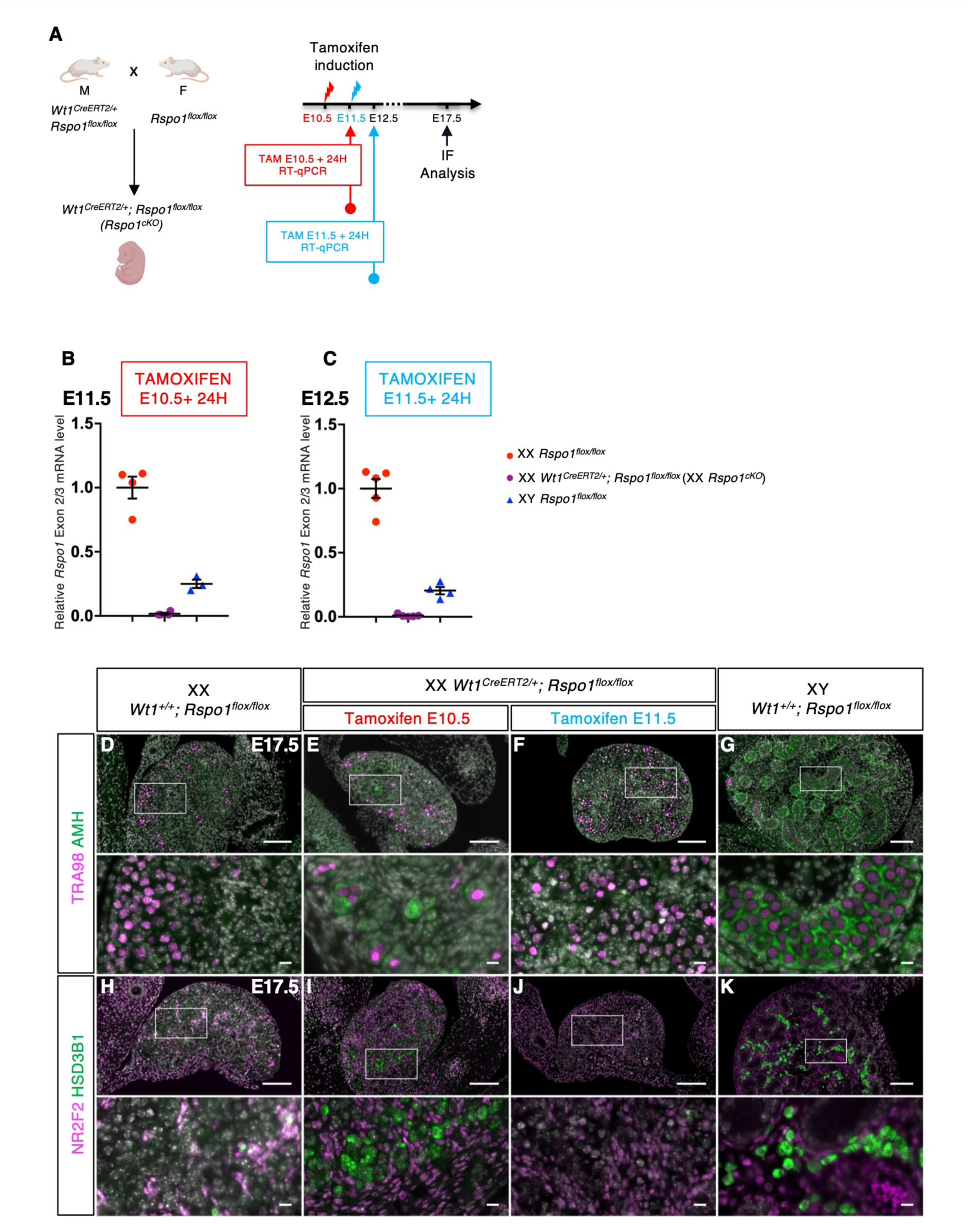
Generation of *Rspo1* conditional mutant in somatic gonadal cells. (**A**) Strategy to delete *Rspo1* exon 2 in somatic gonadal cells using the *Wt1^CreERT2^* line and tamoxifen administration at embryonic day 10.5 (E10.5) or E11.5. (**B**) Quantification of *Rspo1* transcripts containing exon 2 by RT-qPCR after normalization to *Sdha* at E11.5 (24 hours after tamoxifen administration at E10.5). (**C**) Quantification of *Rspo1* transcripts containing exon 2 by RT-qPCR after normalization to *Sdha* at E12.5 (24 hours after tamoxifen administration at E11.5). Data are shown as means ± SEM. (**D** to **G**) Immunofluorescence for the germ cell marker TRA98 (magenta) and the mature granulosa and Sertoli cell marker AMH (green) in the indicated genotypes at E17.5. (**H** to **K**) Immunofluorescence for the interstitial/stromal progenitor marker NR2F2 (magenta) and the steroidogenic cell marker HSD3B1 (green) in the indicated genotypes at E17.5. Nuclei stained with Hoechst 33342 are shown in grey. Data are representative of triplicate biological replicates. Scale bar = 100 µm. Scale bar = 10 µm in zoomed insets.

**Fig. S3.**
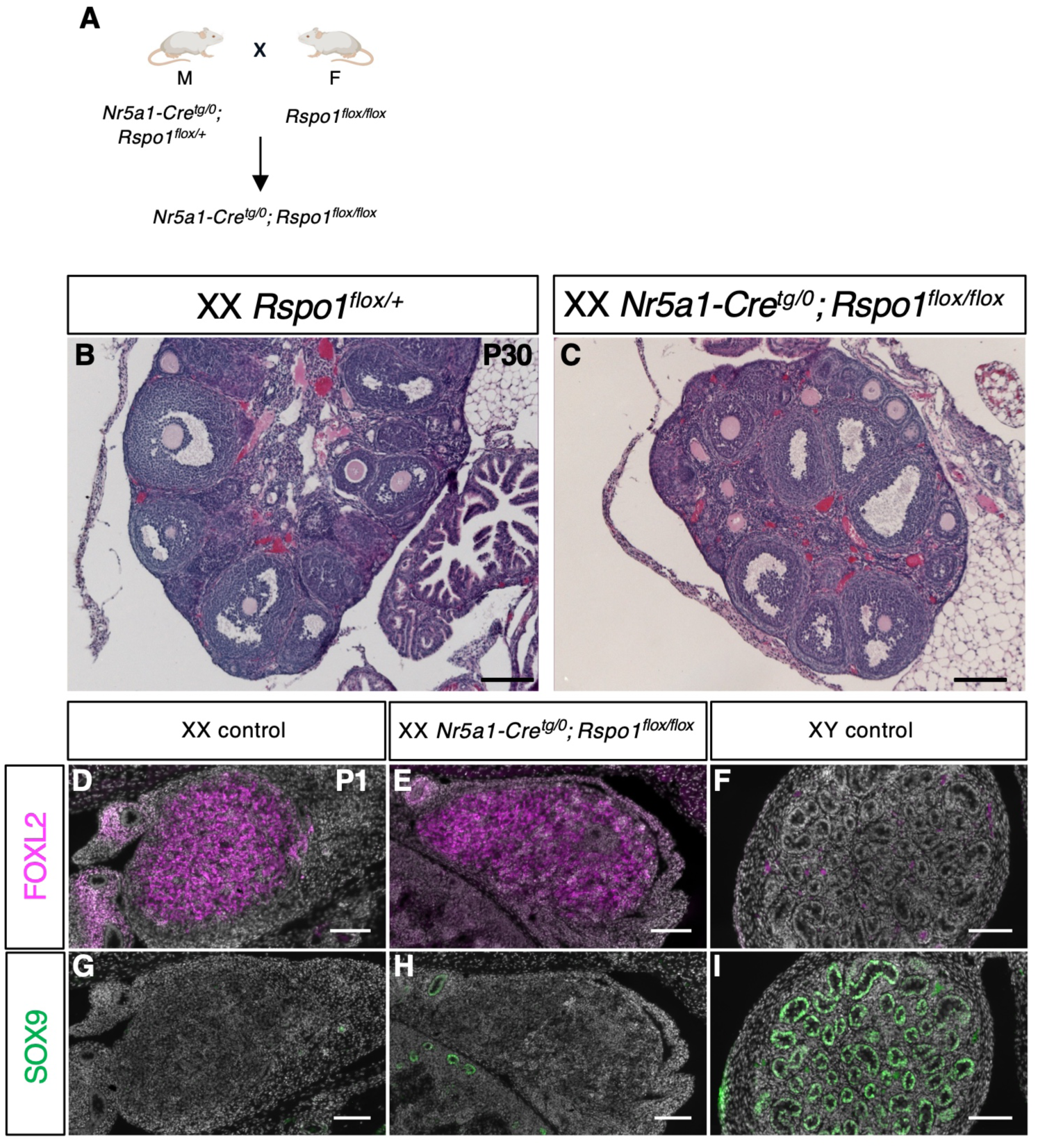
Generation of *Rspo1* conditional mutant in somatic gonadal cells with *Nr5a1-Cre*. (**A**) Strategy to delete *Rspo1* exon 2 in somatic gonadal cells using the *Nr5a1-Cre* line. (**B** and **C**) Hematoxylin/eosin staining of sections of XX *Rspo1^flox/+^* gonads and XX *Nr5a1-Cre; Rspo1^flox/flox^* gonads at post-natal day 30 (P30). Both gonads are ovaries containing developing follicles (arrows). Only 1 out of 16 XX *Nr5a1-Cre; Rspo1^flox/flox^*mutants at P30 showed an ovotestis phenotype. Scale bar = 200 µm. (**D** to **F**) Immunofluorescence for the pre-granulosa cell marker FOXL2 in the indicated genotypes at P1. (**G** to **I**) Immunofluorescence for the Sertoli cell marker SOX9 in the indicated genotypes at P1. Nuclei stained with Hoechst 33342 are shown in grey. Data are representative of triplicate biological replicates. Scale bar = 100 µm.

**Fig. S4.**
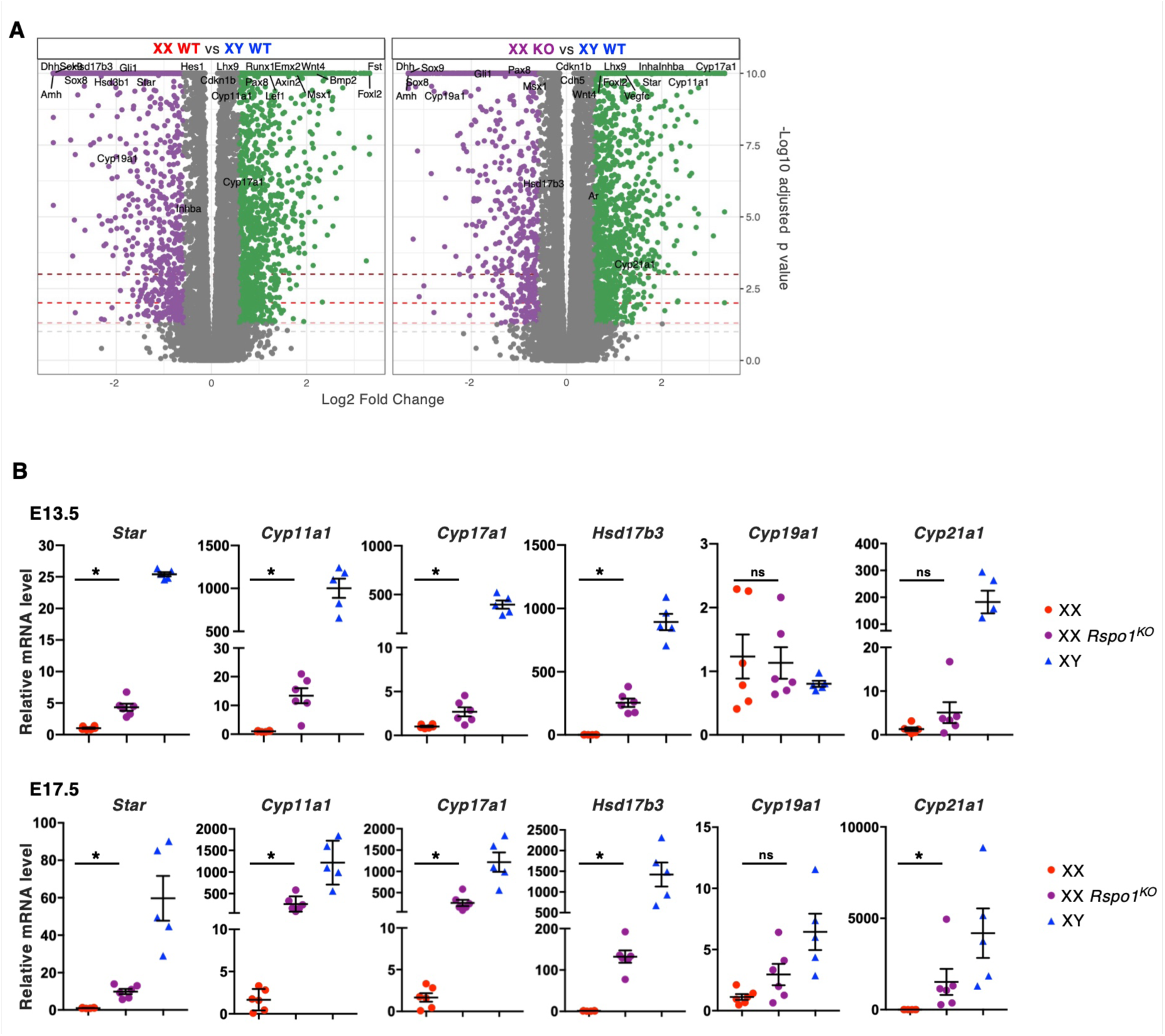
Upregulation of genes related to androgen synthesis in XX *Rspo1^KO^*gonads. (**A**) Volcano plot shown differentially expressed genes between XX WT and XY WT gonads or XX KO and XY WT gonads (-Log10 adjusted p value > 1.25). Downregulated genes (Log2 Fold Change < -0.5, purple) and up-regulated genes (Log2 Fold Change > 0.5, green) are shown. Data from E12.0 bulk RNA-sequencing analysis described in Fig. 2A. (**B**) Quantification of *Star*, *Cyp11a1*, *Cyp17a1*, *Hsd17b3*, *Cyp19a1* and *Cyp21a1* transcripts by RT-qPCR after normalization to *Sdha* and *Tbp* at E13.5 and E17.5. Data are shown as means ± SEM. Statistical significance was assessed by Mann-Whitney U two-tailed test. * indicates P value ≤ 0.05. ns indicates P value > 0.05.

**Fig. S5.**
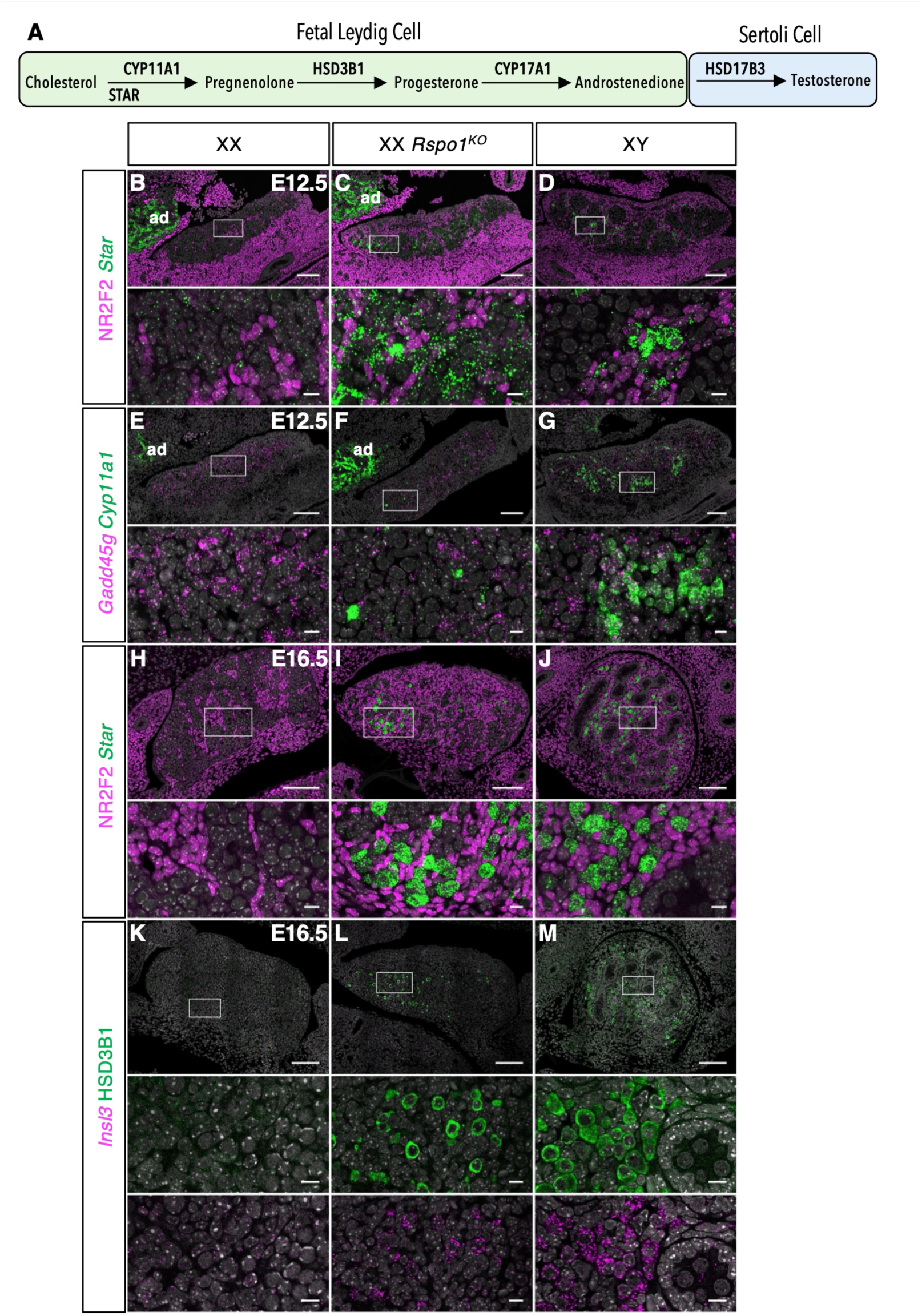
Androgens production in XX *Rspo1* mutant gonads. **(A**) Schematic representation of androgens production in the mouse fetal testis by the joined action of Fetal Leydig cells and Sertoli cells. (**B** to **D**) RNAscope in situ hybridization to detect *Star* transcripts (green) and immunofluorescence to detect NR2F2 (magenta) in the indicated genotypes at E12.5. ad: adrenal gland. (**E** to **G**) RNAscope in situ hybridization to detect *Cyp11a1* (green) and *Gadd45g* (magenta) in the indicated genotypes at E12.5. ad: adrenal gland. (**H** to **J**) RNAscope in situ hybridization to detect *Star* transcripts (green) and immunofluorescence to detect NR2F2 (magenta) in the indicated genotypes at E16.5. (**K** to **M**) RNAscope in situ hybridization to detect *Insl3* transcripts (magenta) and immunofluorescence to detect HSD3B1 (green) in the indicated genotypes at E16.5. Nuclei stained with Hoechst 33342 are shown in grey. Data are representative of triplicate biological replicates. Scale bar = 100 µm. Scale bar = 10 µm in zoomed insets.

**Fig. S6.**
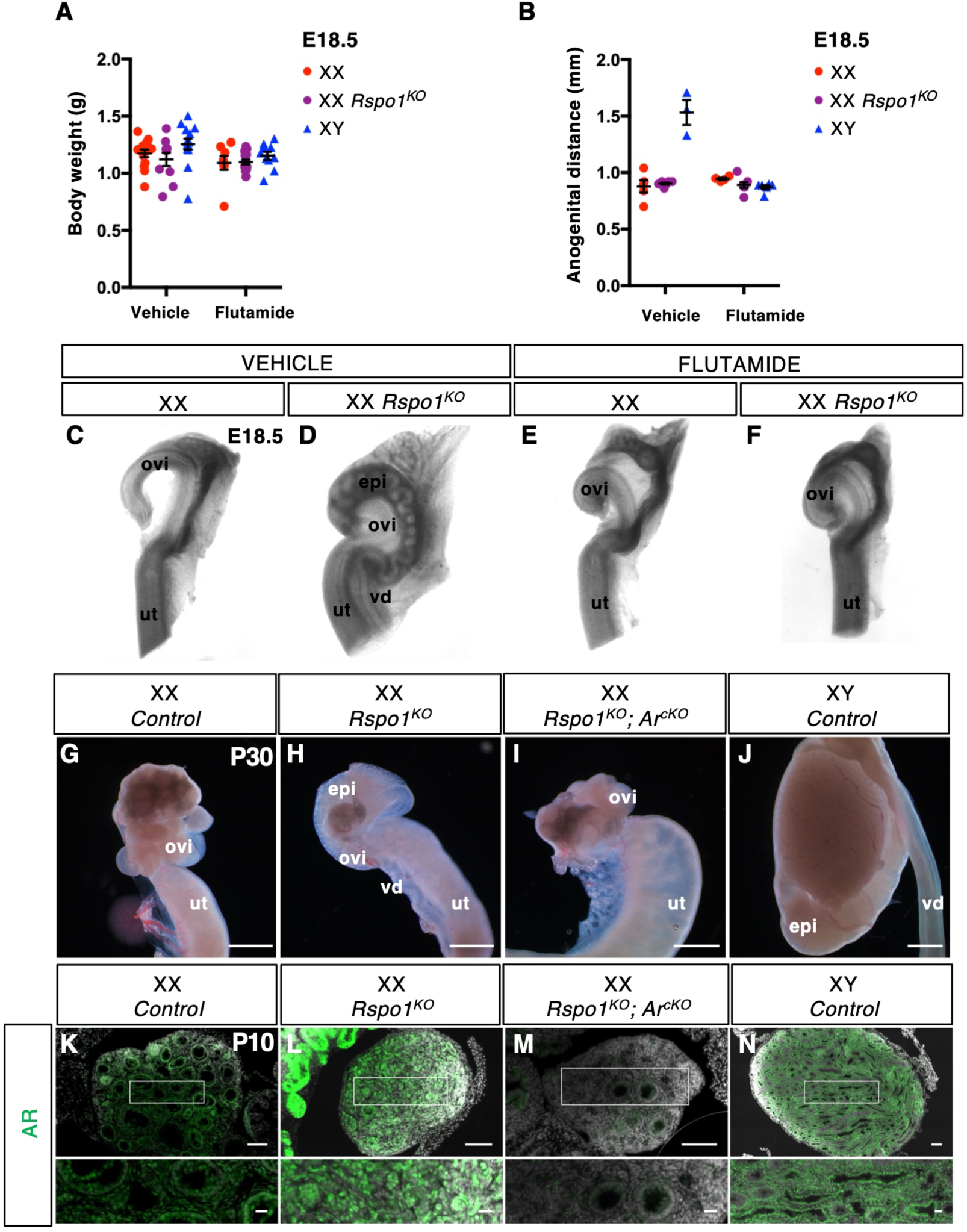
Inhibition of Androgen signaling in XX *Rspo1* mutant gonads. (**A**) Body weight of E18.5 individuals of the indicated genotypes after treatment with vehicle or flutamide of pregnant mothers from E10.5 to E18.5. Data are shown as means ± SEM. (**B**) Anogenital distance measured on E18.5 individuals of the indicated genotypes after treatment with vehicle or flutamide of pregnant mothers from E10.5 to E18.5. Data are shown as means ± SEM. The anogenital distance of XY embryos treated with flutamide is reduced, demonstrating the efficient inhibition of androgen signaling. (**C** to **F**) Macroscopic view of dissected internal genitalia of XX control and *Rspo1^KO^* E18.5 individuals after treatment with vehicle or flutamide of pregnant mothers from E10.5 to E18.5. Vehicle treated XX *Rspo1^KO^* individuals show development of both female (ovi: oviduct, and ut: uterus) and male (epi: epididymis and vd: vas deferens) structures. Flutamide treated XX *Rspo1^KO^* individuals show development of female structures only, demonstrating the efficient inhibition of androgen signaling. (**G** to **J**) Macroscopic view of gonads and internal genitalia of P30 in the indicated genotypes. XX *Rspo1^KO^*individuals show development of both female (ovi: oviduct and ut: uterus) and male (epi: epididymis and vd: vas deferens) structures. XX *Rspo1^KO^;Ar^cKO^* individuals show development of female structures only. Scale bar = 100 µm. (**K** to **N**) immunofluorescence staining for AR (green) on gonadal sections at P10 in the indicated genotypes. Nuclei stained with Hoechst 33342 are shown in grey. Data are representative of triplicate biological replicates. Scale bar = 100 µm. Scale bar = 20 µm in zoomed insets.

**Table S1.**
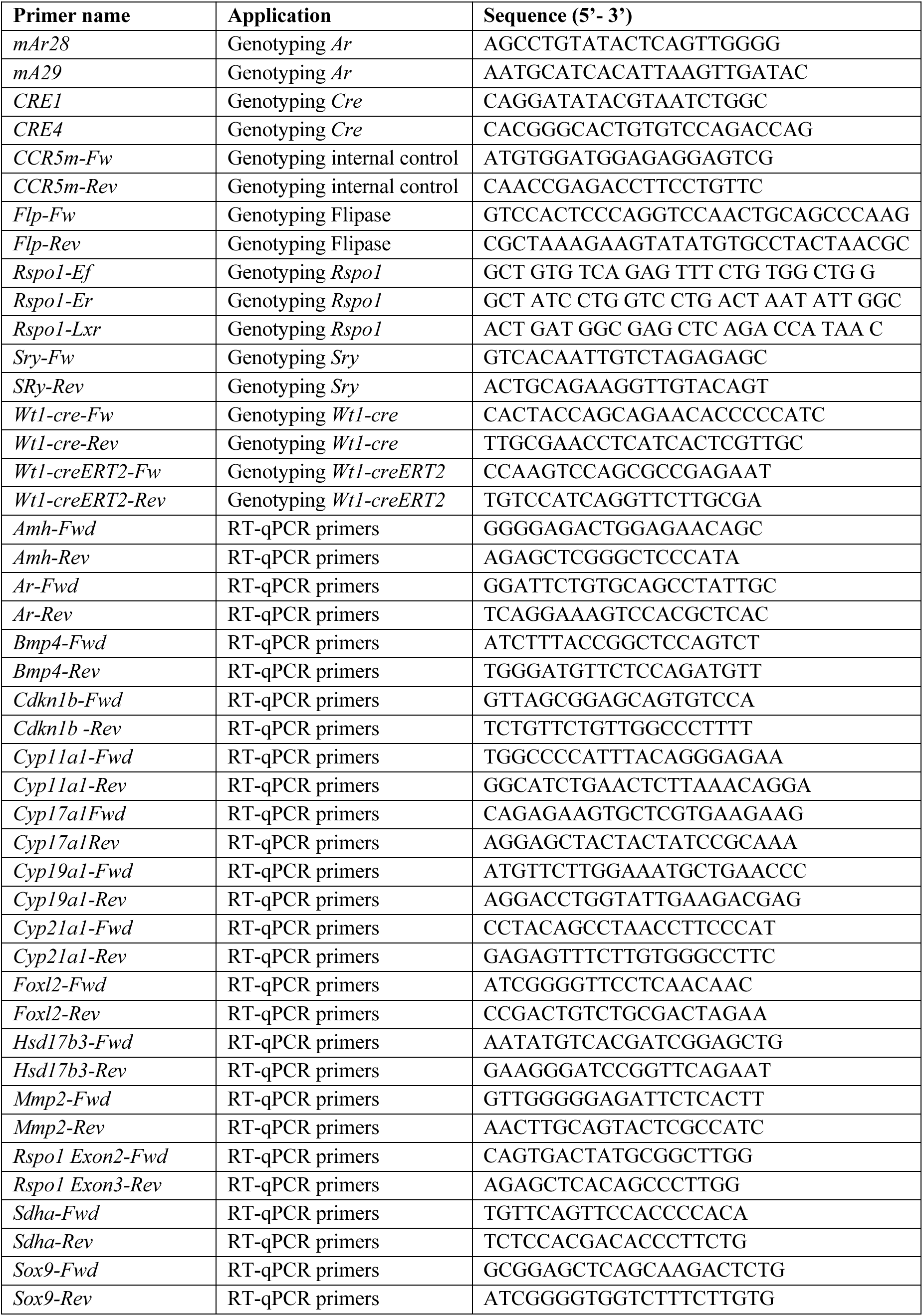
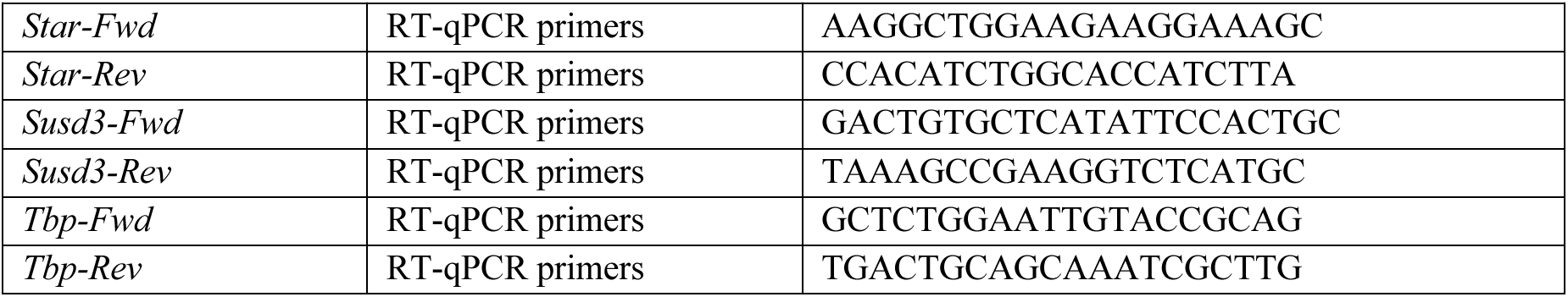
Primers used in this study.

**Table S2.**
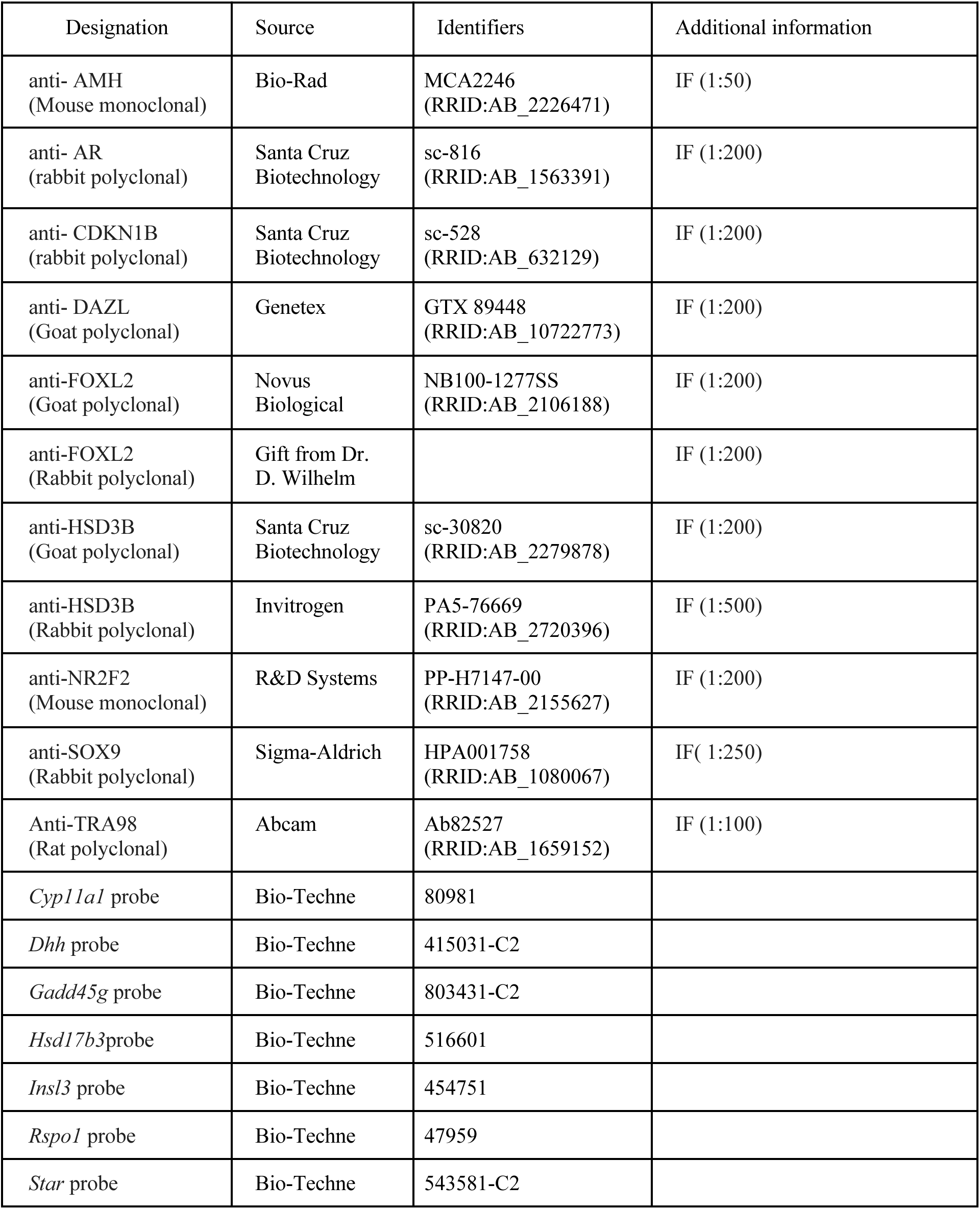
Antibodies and probes used in this study.

**Table S3.**
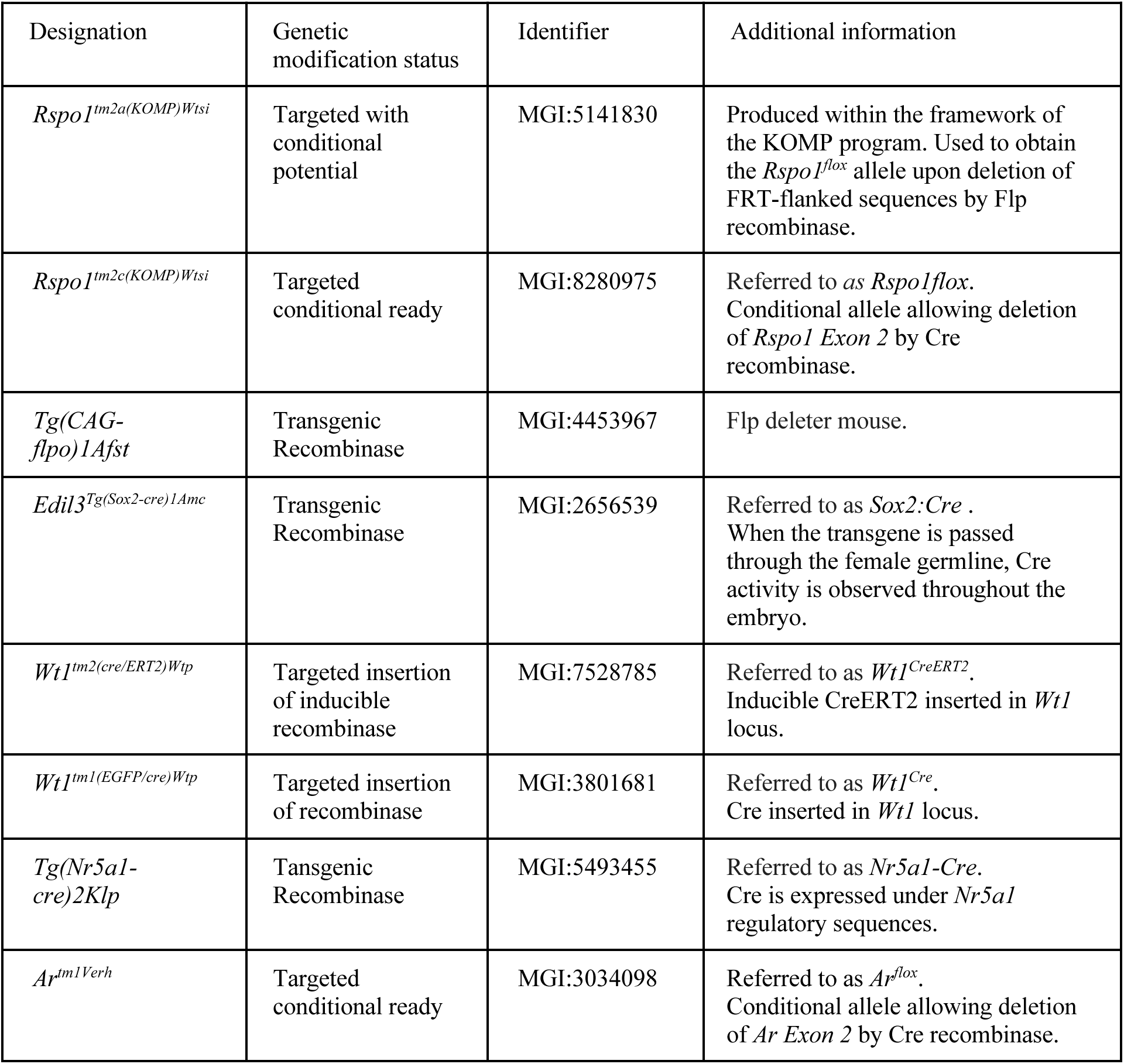
Mouse lines used in this study.

